# Nitrate-driven dark oxygen production by diverse deep-sea microorganisms

**DOI:** 10.1101/2025.03.10.642362

**Authors:** Rikuan Zheng, Chong Wang, Hongliang Wang, Haibin Qi, Chaomin Sun

## Abstract

To date, only three biotic pathways for light-independent oxygen production (i.e., dark oxygen) have been reported: chlorate dismutation, nitric oxide dismutation, and methanobactin-dependent water lysis. Oxygen has been shown to be produced and consumed in dark and anoxic environments, as evidenced by its prevalence in deep-sea surface sediments. However, the microbial communities and pathways driving deep-sea dark oxygen production (DOP) remain poorly understood. Here we identify a novel DOP pathway driven by dissimilatory nitrate reduction to ammonium (DNRA) in two deep-sea *Deferribacterota* strains. The DOP activity of these *Deferribacterota* strains potentially promotes the formation of manganese nodules and influences the growth and metabolism of both aerobic and anaerobic microbes, underscoring the significant role of microbial DOP in shaping deep-sea geological and ecological systems. We also cultured several deep-sea DOP-capable microorganisms from the *Campylobacterota*, *Deinococcota*, and *Pseudomonadota* phyla, which generate dark oxygen through unidentified nitrate-driven pathways. Combined with previous geological evidence, our results suggest that nitrate-driven dark oxygen, alongside photosynthetic oxygen, jointly contributed to early Earth’s oxygenation, driving the Great Oxidation Event. Overall, our findings provide novel insights into deep-sea oxygen sources, shed light on the origins of early Earth’s oxygen, and expand perspectives on the potential for terraforming other planets.

---

Photosynthetic oxygen is widely regarded as the primary source of oxygen on Earth (*1, 2*). Light-independent abiotic or biotic process also contributes to oxygen production, particularly during early Earth history (the oxygen-free age), a process termed dark oxygen production (DOP) (*3*). Recent studies have suggested that dark oxygen-generating microorganisms may play a significant role in certain anoxic environments (*4*). Microbial DOP has not been extensively studied, with only three known pathways: chlorite (ClO_2_^-^) dismutation (ClO_2_^-^ → Cl^-^ + O_2_) (*5*), nitric oxide (NO) dismutation (2NO_2_^-^ → 2NO → N_2_ + O_2_ or 4NO_2_^-^ → 4NO → 2N_2_O + O_2_) (*6, 7*), and water lysis by methanobactins (*8*). In addition, reactive oxygen species (e.g., H_2_O_2_) can be disproportionated to O_2_ and H_2_O both biotically, via enzymes such as superoxide dismutase and catalase, and abiotically, by ferrous iron and other reduced metals (*9, 10*). However, due to the low levels of reactive oxygen species *in vivo*, the resulting oxygen production is negligible and difficult to detect, rendering it insignificant compared to the other three biotic DOP pathways, particularly in most low-temperature, anoxic systems (*3*).

Growing geochemical and microbiological evidences suggest that DOP is a significant and widespread, yet often overlooked, process in nominally anoxic environments, including groundwater, subsurface, lake, wetland, and marine ecosystems. Groundwater and subsurface environments harbor strictly aerobic methanotrophs (*11*), methanotrophic *Methylobacter* (*4*), and other viable aerobic heterotrophs (*4*), indicative of relatively oxidizing conditions (*12*). Aerobic methanotrophs have also been detected in anoxic sediments of Lake Constance, Germany (*13*) and at a methane seep in Lake Qalluuraq, Alaska (*14*), where they are present in sediments up to 70 cm depth. Similarly, an aerobically methanotrophic community has been observed in an Arctic wetland, exhibiting peak activity in the anoxic and transition zones (*15*). Aerobic activity has also been detected several meters below the seafloor within the permanently anoxic Arabian Sea oxygen minimum zone (OMZ), where *Nitrosopumilus* archaeon (*7*) (potential dark oxygen producers), *nod* genes (*6*) (encoding nitric oxide dismutase for DOP), and transcribed oxidase genes were identified (*16*). Moreover, in deep-sea methane seeps and mud volcanoes, aerobic methanotrophs are abundant in methane-rich sediments well below the measurable oxygen penetration depth (*17, 18*). Not only microorganisms, animals (e.g., blind shrimp and *Alvinocaris* shrimp around deep-sea hydrothermal vents; squat lobsters and mussels around cold seeps) also widely distribute in the deep sea, indicating substantial oxygen availability in these regions. What remains intriguing is the origin of deep-sea oxygen. Recent findings suggest that polymetallic nodules at the abyssal seafloor may produce dark oxygen through seawater electrolysis, although the mechanistic details remain unknown (*19*). This observation highlights the potential importance of DOP in deep-sea ecosystems and its implications for generating oxygenated habitats on other ocean worlds. Given the prevalence of oxygen in deep-sea surface sediments (*19*), we are intrigued to explore whether and how microbial DOP contributes to the deep-sea oxygen source.

## Deep-sea *Deferribacterota* bacteria produce dark oxygen

To enrich deep-sea microorganisms capable of producing dark oxygen, we specifically employed a nitrate-containing medium, as denitrification potentially drives microbial DOP (*6, 7, 20*) and nitrate levels are closely correlated with deep-sea oxygen content (*21*). The enrichment was incubated under anaerobic conditions, and the oxygen indicator resazurin—whose reduced form, dihydroresazurin, transitions from colorless to pink when oxidized to resorufin (fig. S1A)—was added to the medium to detect the presence of trace oxygen. Surprisingly, we observed that two anaerobic tubes supplemented with hydrothermal vent samples exhibited visible microbial growth and transitioned from colorless to pink (fig. S1B), suggesting that these two enrichments might produce oxygen. Two rod-shaped bacterial strains (DY0037 and DY0609) were isolated and purified from the two enrichments (fig. S1C) and identified as members of the *Deferribacterota* phylum (fig. S1D). To further confirm oxygen production by *Deferribacterota* strains DY0037 and DY0609, we conducted scaled-up incubations in larger vessels. Over time, we observed the formation of gas bubbles in the anaerobic bottles (fig. S2A), accompanied by a gradual color change in the medium from colorless to pink (24 hours) and ultimately to deep red (48 hours) (Fig. 1A). This phenomenon was further supported by measurements of dissolved oxygen using a liquid phase oxygen electrode. These analyses revealed a gradual increase in dissolved oxygen concentrations with culture growth, reaching approximately 70 μM (Fig. 1, B and C). Subsequently, strain DY0037 exhibited a decline in oxygen levels, indicating consumption, while strain DY0609 maintained a stable dissolved oxygen concentration, likely due to its low oxygen-utilization capacity. The dissolved oxygen concentration is over 300-fold higher than that produced by a previously reported ammonia-oxidizing archaea via NO dismutation (*7*), and nearly exceeds the maximum dissolved oxygen level achievable in anoxic aquatic habitats (<60 μM) (*22*), as well as the upper limit of typical marine hypoxic conditions (2 mg/L or 63 μM) (*23*).

**fig. S1.**
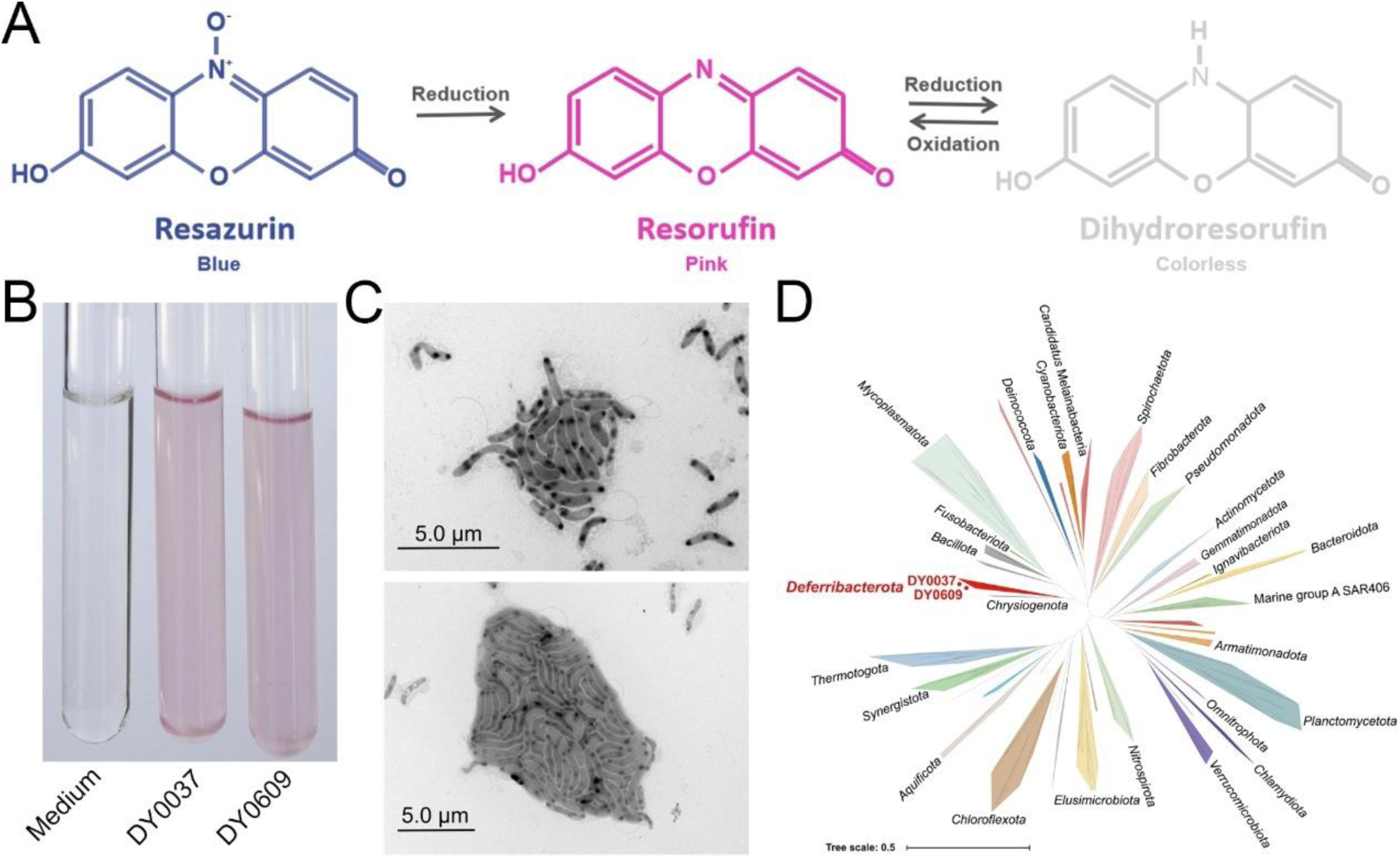
Isolation, morphology and phylogenetic affiliation of two deep-sea anaerobic bacteria with potential oxygen-production capability. (**A**) The oxygen indicator resazurin exhibits distinct color states depending on oxygen availability: blue (oxidized form, resazurin), pink (intermediate reduced form, resorufin), and colorless (fully reduced form, dihydroresorufin). (**B**) Observation of color changes in the control medium compared to the same medium inoculated with deep-sea bacterial strains DY0037 or DY0609. (**C**) Transmission electron micrographs showing the morphology of strains DY0037 and DY0609. Bar is 5 μm. (**D**) Phylogenetic analysis of strains DY0037 and DY0609 based on full-length 16S rRNA gene sequences from some *Deferribacterota* members and other representative phyla. Scale bar, 0.5 substitutions per nucleotide position.

**Fig. 1.**
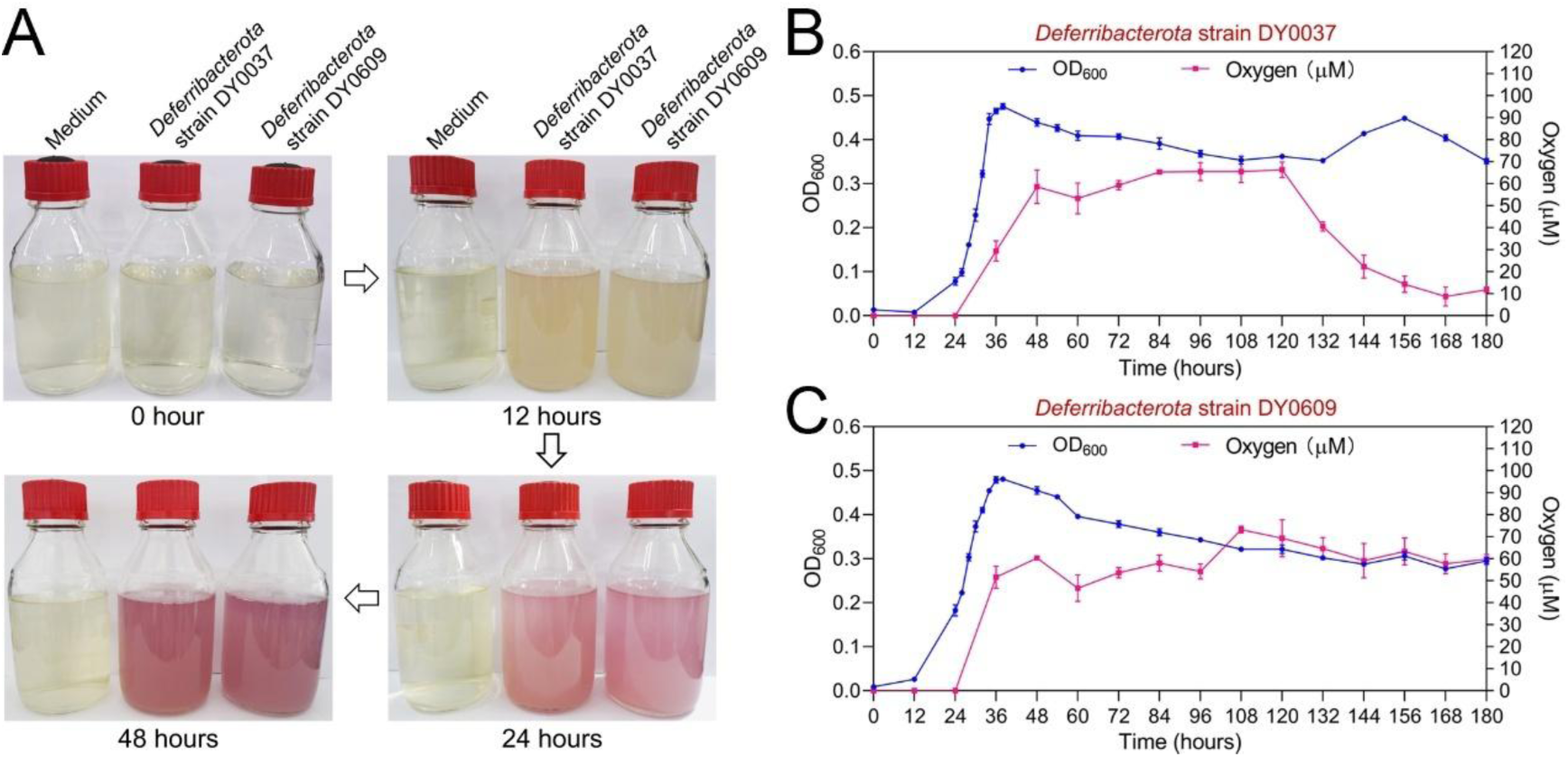
Deep-sea *Deferribacterota* strains DY0037 and DY0609 produce oxygen during anaerobic growth. (**A**) Time-lapse images showing the culture medium transitioning from colorless (0 hour) to slightly pink (12 hours), pink (24 hours) and finally red (48 hours) during anaerobic growth of strains DY0037 and DY0609. (**B** and **C**) Growth curves and dissolved oxygen production profiles of strains DY0037 (B) and DY0609 (C).

It is noteworthy that the growth and oxygen production of the two *Deferribacterota* strains are strictly dependent on nitrate (fig. S2B). In contrast, no growth or oxygen production was observed when cells were killed by high-temperature, high-pressure sterilization (fig. S3), ruling out the possibility of abiotic oxygen production or oxygen intrusion into the incubation vessels. Furthermore, incubations of strains DY0037 and DY0609 in above nitrate-containing medium, with or without pyruvate supplementation, showed no significant difference in growth (fig. S4A) and dissolved oxygen production (fig. S4B). This result excluded oxygen production via H_2_O_2_ dismutation (H_2_O_2_ → H_2_ + O_2_), as pyruvate abiotically scavenges H_2_O_2_ through decarboxylation (*24*). The absence of chlorate and chlorite in the culture medium and throughout the cultivation process also excluded the possibility of chlorite-dependent oxygen production. In addition, incubation of *Deferribacterota* strains in nitrate-containing medium revealed no significant difference in growth (fig. S4C) or dissolved oxygen production (fig. S4D) under dark and light conditions, confirming these two *Deferribacterota* strains produce dark oxygen. *Deferribacterota* bacteria are prevalent in deep-sea habitats (*25–27*), and are able to oxidize complex organic compounds and organic acids (mainly acetate) using a variety of electron acceptors (mainly nitrate) (*28*). Therefore, nitrate-dependent oxygen production may be widespread among *Deferribacterota* species, potentially contributing to oxygen availability across diverse deep-sea ecosystems.

**fig. S2.**
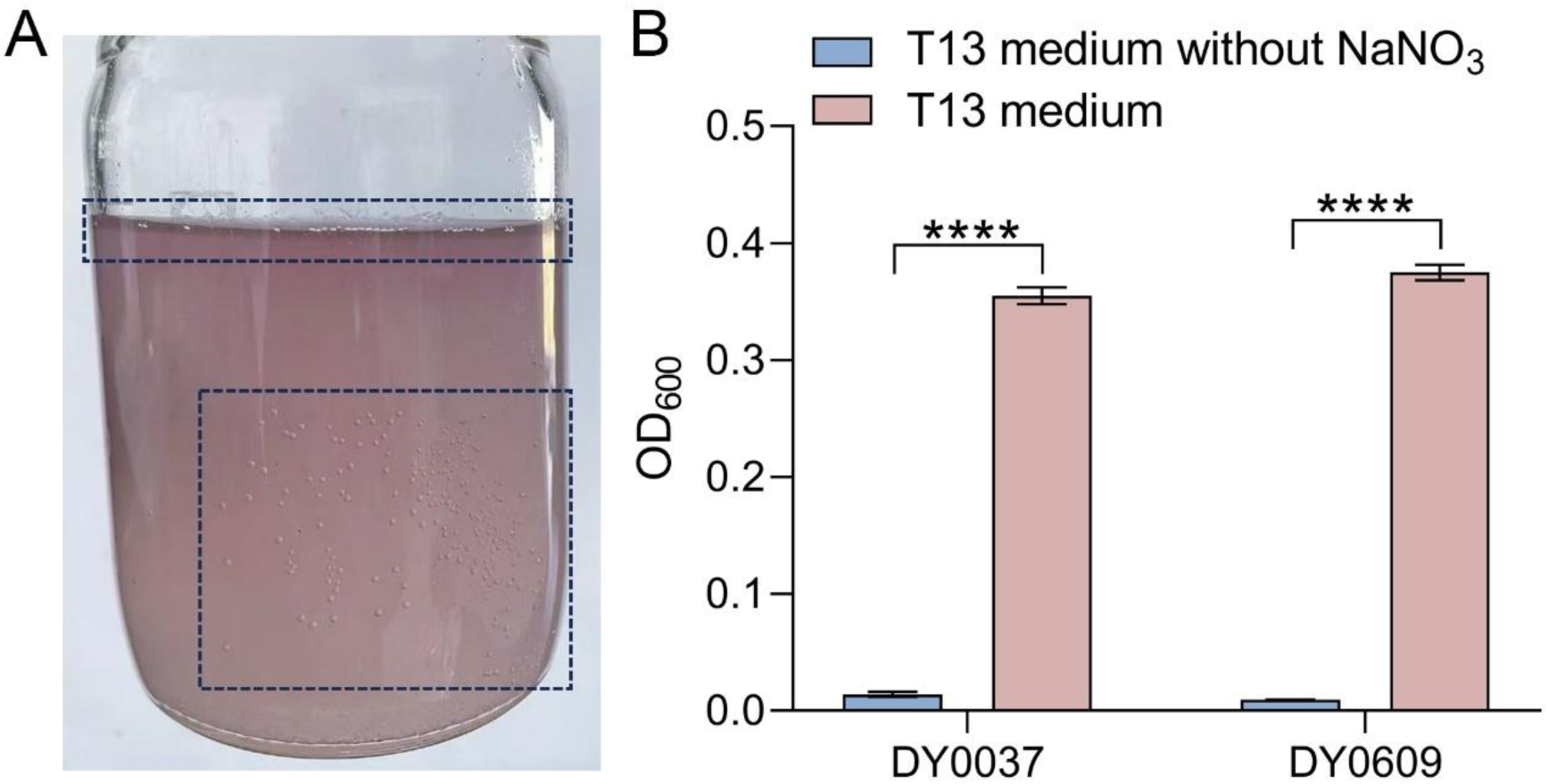
*Deferribacterota* strains DY0037 and DY0609 produce oxygen strictly dependent on the presence of nitrate. (**A**) Oxygen production is apparent on the inner walls and in the upper layer of the culture bottles (dashed boxes) during anaerobic cultivation of strains DY0037 and DY0609 in the nitrate-containing medium. (**B**) Growth profiles of strains DY0037 and DY0609 in T13 medium supplemented with and without sodium nitrate. Error bars show mean ± SD. \*\*\*\**P* < 0.0001.

**fig. S3.**
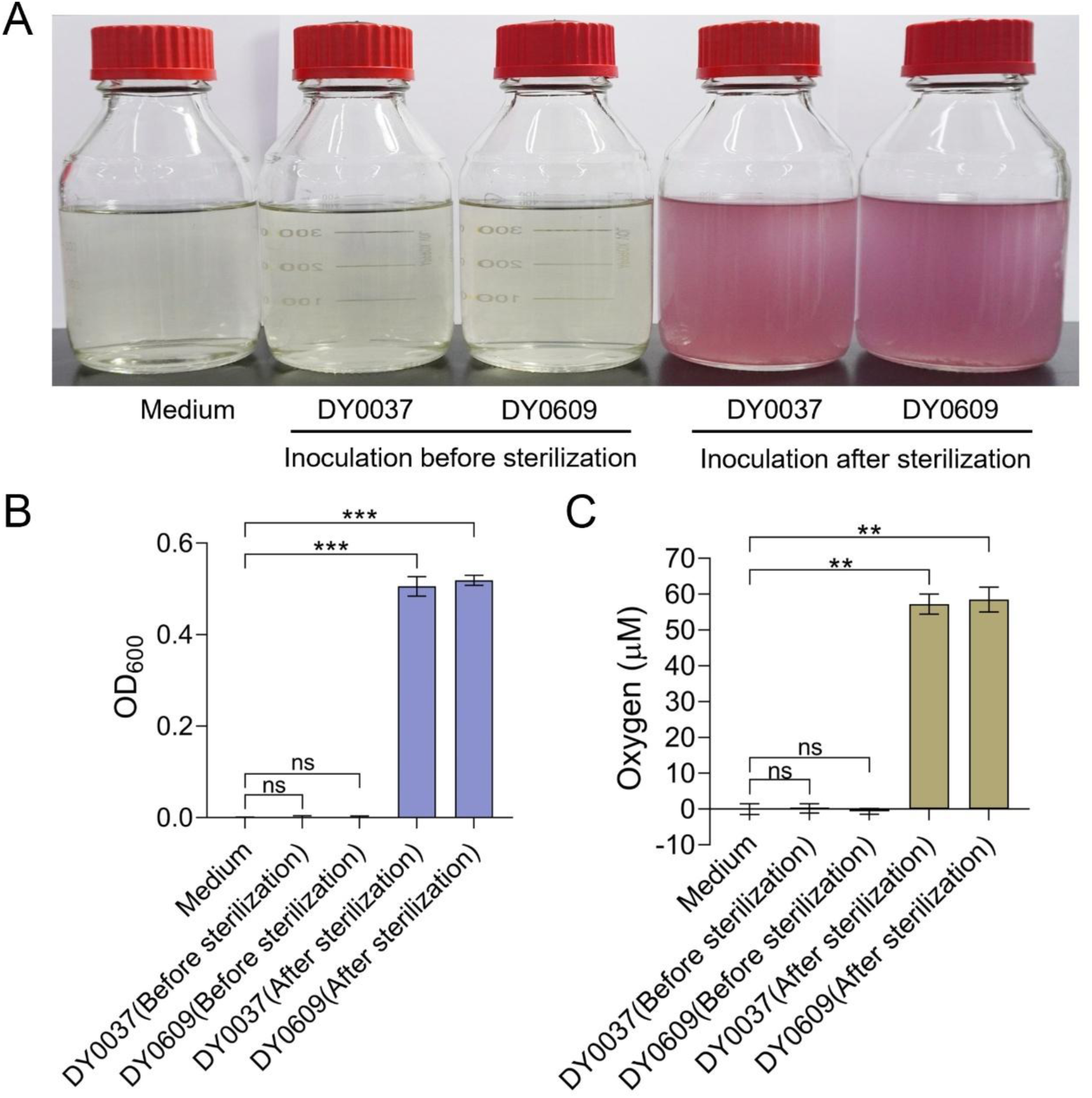
Ruling out the possibility of abiotic oxygen production and oxygen intrusion into the incubation vessels during cultivation of *Deferribacterota* strains DY0037 and DY0609. (**A**) Photographs of culture medium after adding seed cultures of strains DY0037 and DY0609, both before and after sterilization, with uninoculated medium serving as the control. (**B**) Growth profiles of strains DY0037 and DY0609 following the addition of seed cultures before and after medium sterilization, compared to the control. (**C**) Dissolved oxygen levels measured following the addition of seed cultures of strains DY0037 and DY0609 before and after medium sterilization, with uninoculated medium serving as the control. Error bars show mean ± SD. \*\**P* < 0.01, \*\*\**P* < 0.001.

**fig. S4.**
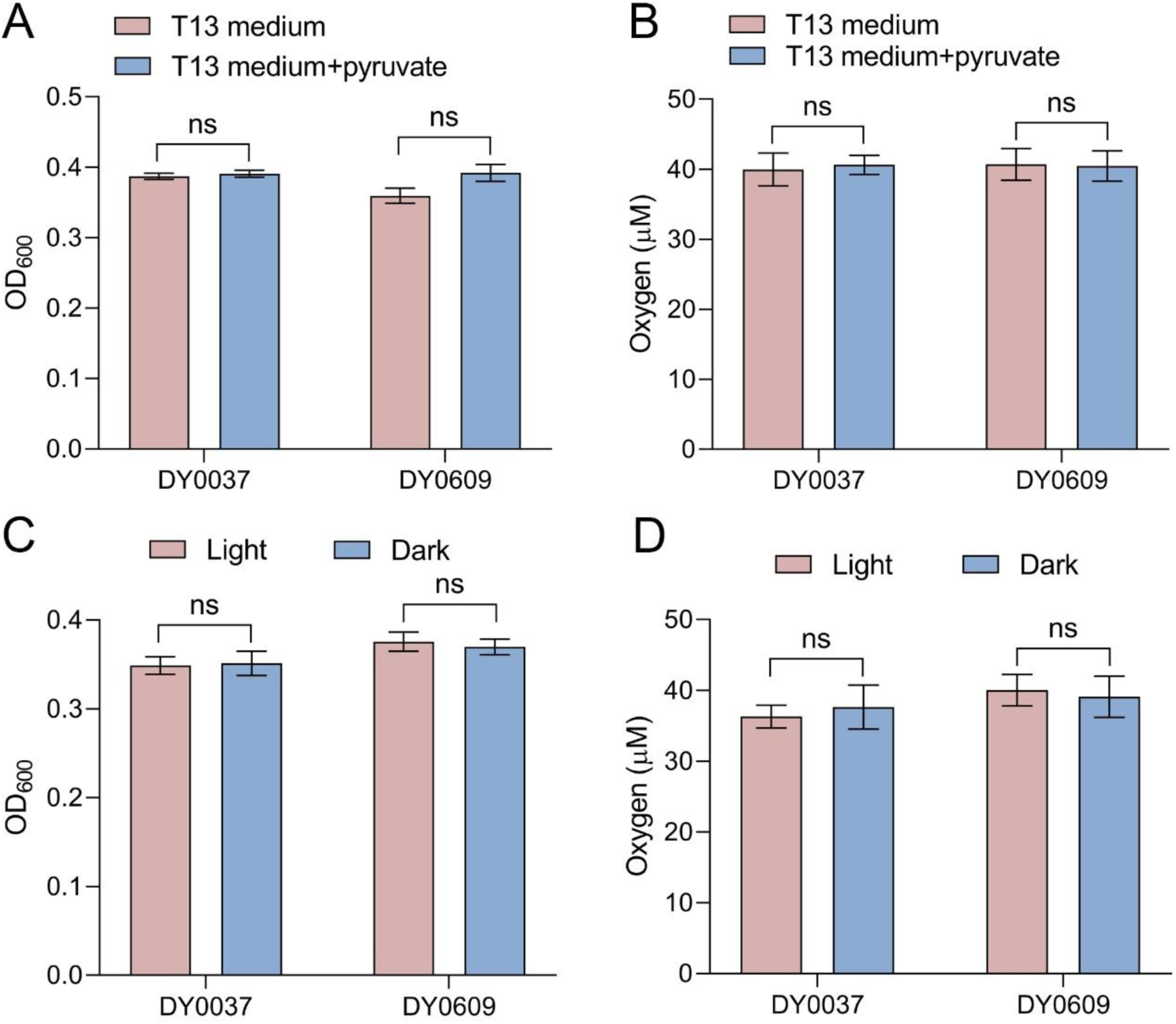
Ruling out the possibility of oxygen production via H_2_O_2_ dismutation in *Deferribacterota* strains DY0037 and DY0609. (**A**) Growth profiles of strains DY0037 and DY0609 in T13 medium with or without 0.2 mM pyruvate supplementation. (**B**) Dissolved oxygen concentrations of strains DY0037 and DY0609 in T13 medium with or without 0.2 mM pyruvate supplementation. (**C**) Growth profiles of strains DY0037 and DY0609 cultivated under light and dark conditions. (**D**) Dissolved oxygen concentrations of strains DY0037 and DY0609 cultivated under light and dark conditions. Error bars show mean ± SD. “ns” denotes no significant difference (*P* > 0.05).

## *Deferribacterota* strains produce dark oxygen through a novel pathway

To elucidate the DOP pathway in *Deferribacterota* strains DY0037 and DY0609, we first sequenced their genomes and analyzed their nitrate reduction pathways, as nitrate is necessary for growth and oxygen production of these two strains. Intriguingly, we found they both possess incomplete dissimilatory nitrate reduction to ammonia (DNRA) and denitrification pathways; specifically, they encode only nitrate reductases (NapADGH and NarGHIJK) and nitrous oxide reductase (NosZ) (Fig. 2A). Cultures grown with ^18^O-nitrate exhibited significantly elevated ^18^O_2_/^16^O_2_ ratios compared to control cultures (7.4-fold and 12.9-fold increases for strains DY0037 and DY0609, respectively), confirming that both strains utilize nitrate for oxygen generation (Fig. 2B). Furthermore, the formation of abundant magnesium ammonium phosphate mineral particles (MgNH_4_PO_4_·6H_2_O) was observed during cultivation, strongly indicating significant ammonium production (fig. S5). This observation, combined with the detection of ^15^N-nitrate, ^15^N-nitrite, and ^15^N-ammonium accumulation (approximately 1.2, 2.1, and 1.5 mM for strain DY0037 and 1.1, 2.0, and 2.0 mM for strain DY0609, respectively) in cultures amended with ^15^N-nitrate (Fig. 2C), as well as the consumption of ammonium during MgNH₄PO₄·6H₂O precipitation, strongly suggests that both strains perform DNRA process. This is despite the absence of genes encoding the enzyme responsible for catalyzing the reduction of NO₂⁻ to NH₄⁺ in their genomes. Real-time monitoring of inorganic nitrogen concentrations during culture growth revealed a significant increase in both ammonium and dissolved oxygen levels, alongside a corresponding decrease in nitrate concentrations. Notably, the total inorganic nitrogen content remained balanced (Fig. 2, D and E), indicating no production of nitrogen-containing gases. Consistent with these findings, transcriptomic analyses under high nitrate stress revealed significant upregulation of genes encoding key enzymes involved in nitrate reduction (fig. S6). Collectively, these results demonstrate that strains DY0037 and DY0609 are capable of performing the DNRA process under anaerobic conditions, leading to oxygen production. Consequently, we propose a novel DNRA-dependent oxygen production pathway in *Deferribacterota* strains DY0037 and DY0609, as illustrated in Figure 2F. This pathway is thermodynamically feasible and distinct from previously reported NO dismutation during denitrification (*6, 7*). In this pathway, after the reduction of nitrate to nitrite by nitrate reductase, an unidentified enzyme facilitates the conversion of one molecule of nitrite into one molecule of oxygen and one molecule of ammonium. This process involves the transfer of two electrons, which may partially originate from intra- or extracellular organic matter (such as sodium acetate), and four protons, which may partly derive from water. Here, we present evidence for a fourth biotic pathway for dark oxygen production (DOP), which could hold significant biological and geochemical importance. Given that *Deferribacterota* strains DY0037 and DY0609 lack canonical nitrite reductases, we propose the existence of a novel enzyme mediating the reduction of nitrite to ammonium and O₂. However, the gene responsible for this process remains unidentified. Future research will focus on developing a genetic system in anaerobic *Deferribacterota* bacteria to identify the key genes controlling dark oxygen generation.

**fig. S5.**
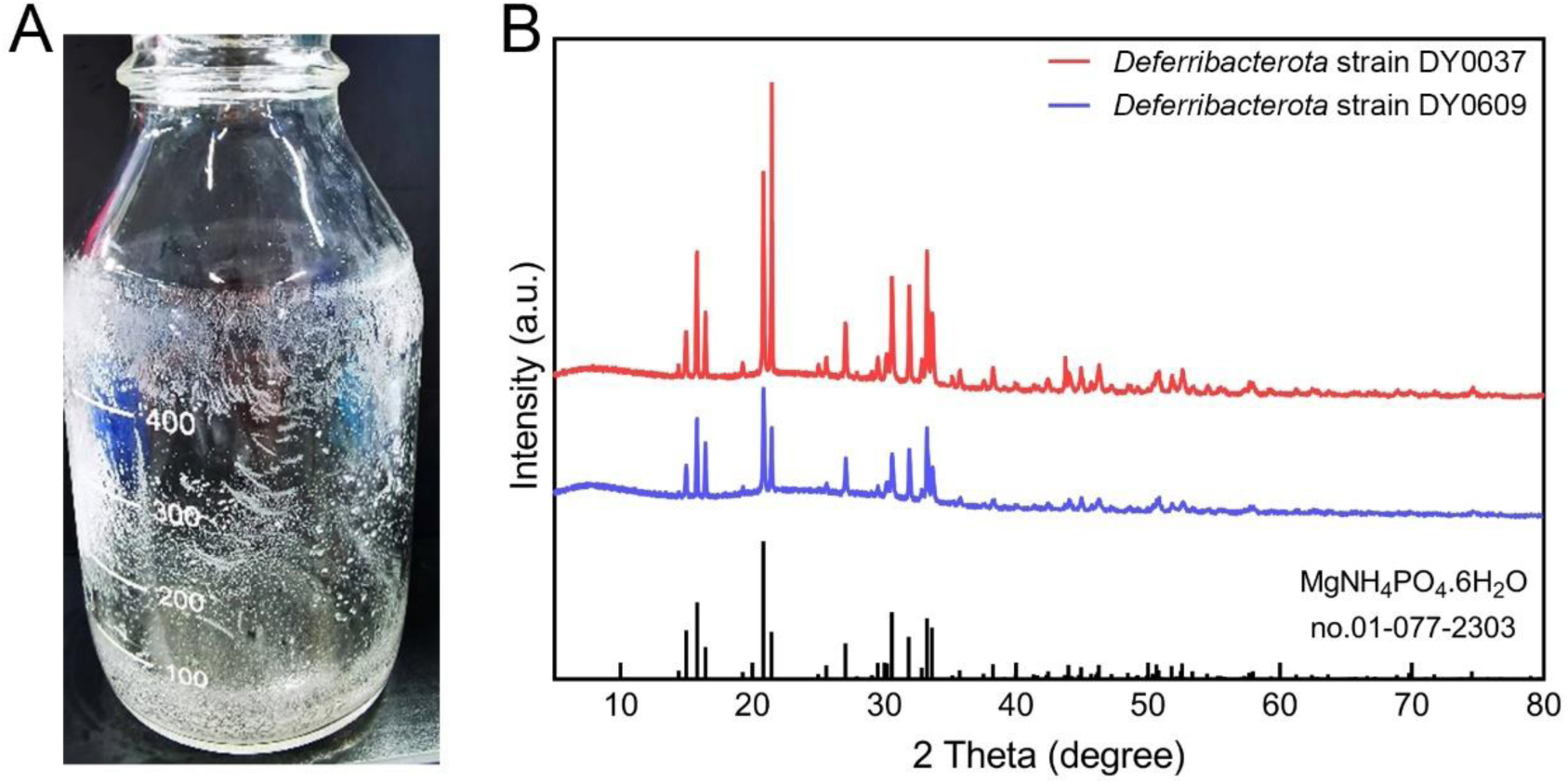
Characterization of minerals formed by *Deferribacterota* strains DY0037 and DY0609 during oxygen production in the nitrate-containing medium. (**A**) Visual observation of abundant mineral formation during the cultivation of strains DY0037 and DY0609. (**B**) X-ray diffraction (XRD) analysis of the minerals formed by strains DY0037 and DY0609.

**fig. S6.**
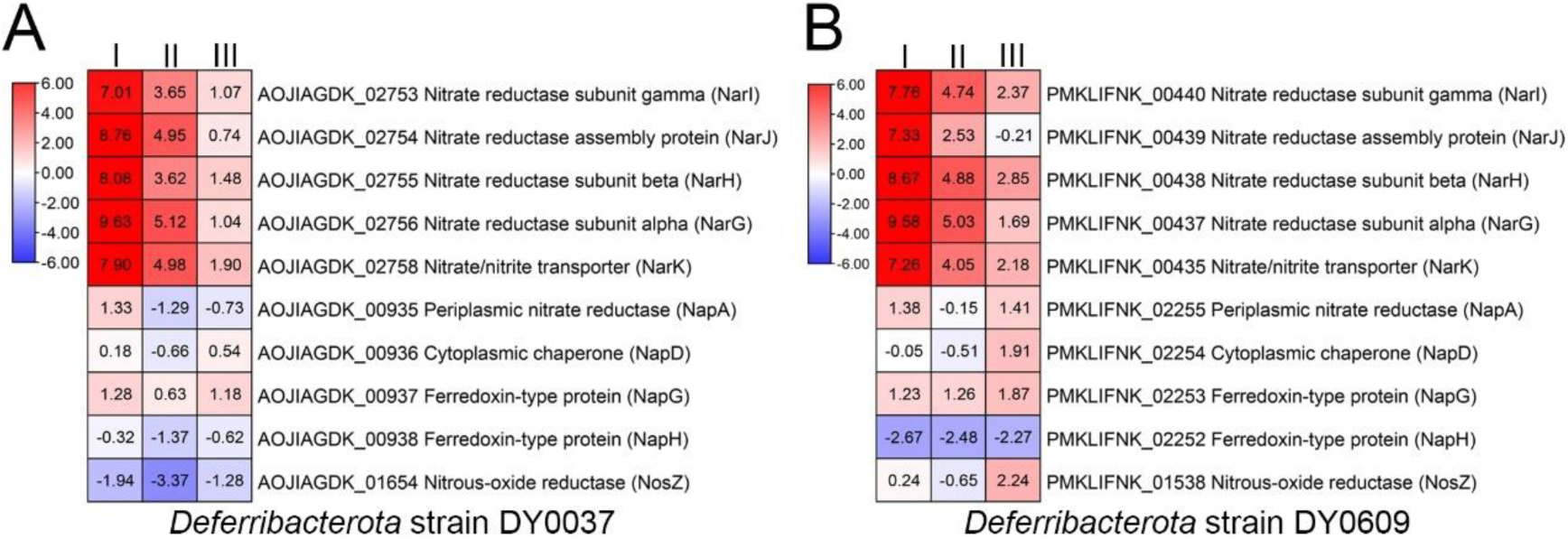
Transcriptomic analysis of nitrogen metabolism-associated gene expression in *Deferribacterota* strains DY0037 and DY0609 in medium containing varying concentrations of nitrate. Transcriptomic profiles of nitrogen metabolism-associated genes in strains DY0037 (**A**) and DY0609 (**B**) under varying nitrate concentrations (0.1, 0.5, 1.0, and 3.0 g/L). “I” represents the ratio of gene expression in strain DY0037 or DY0609 grown in medium with 0.5 g/L nitrate versus 0.1 g/L nitrate; “II” represents the ratio of gene expression in strain DY0037 or DY0609 grown in medium with 1.0 g/L nitrate versus 0.1 g/L nitrate; “III” represents the ratio of gene expression in strain DY0037 or DY0609 grown in medium with 3.0 g/L nitrate versus 0.1 g/L nitrate. The numbers in panels A and B indicate the fold change in gene expression, shown as log_2_ values.

**Fig. 2.**
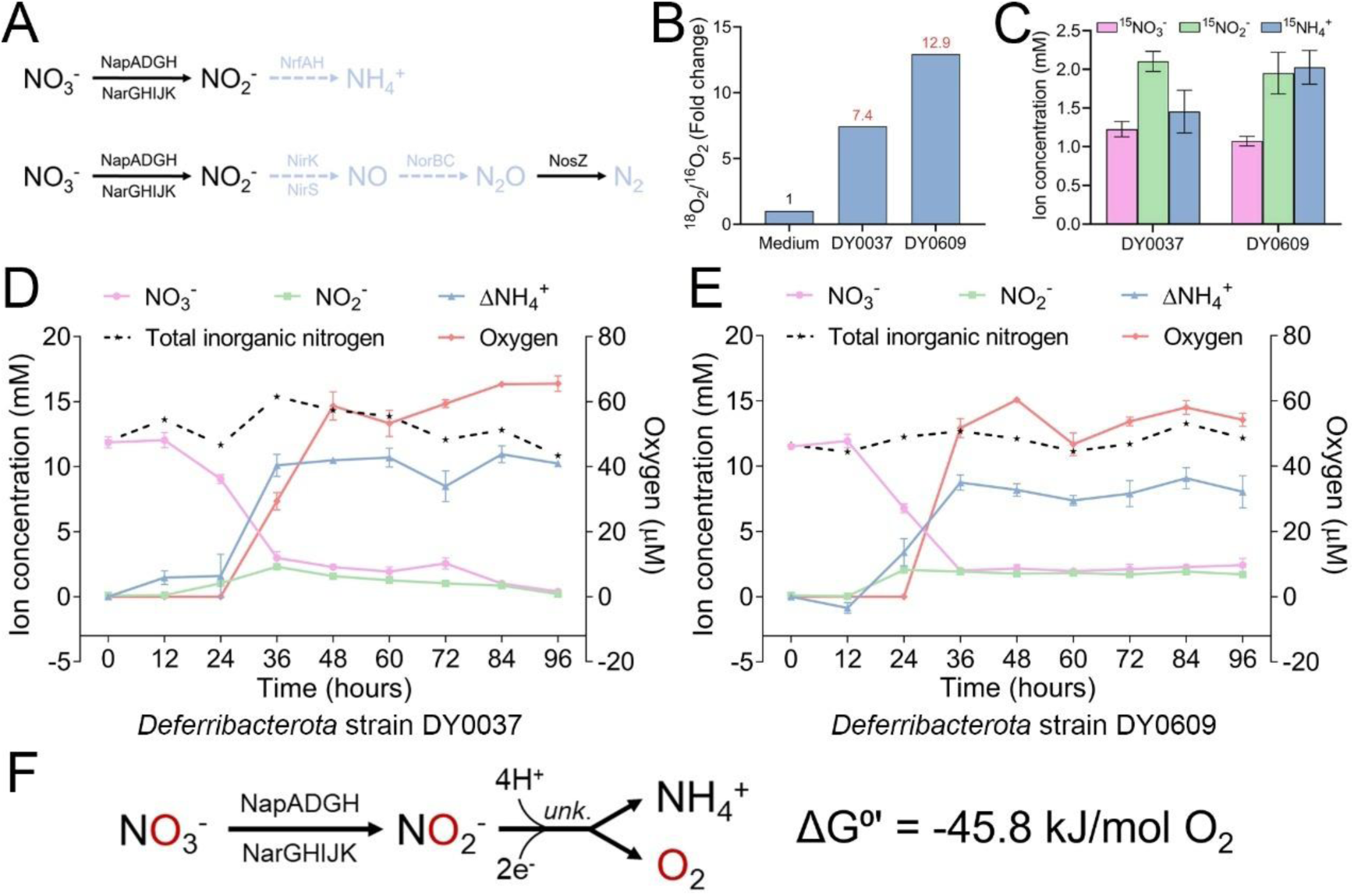
*Deferribacterota* strains DY0037 and DY0609 produce dark oxygen through a DNRA-dependent pathway. (**A**) Genomic analysis of the DNRA and denitrification pathways in *Deferribacterota* strains DY0037 and DY0609. Solid black lines indicate the presence of relevant genes and intermediates, while dashed light blue lines denote the absence of relevant genes and intermediates. (**B**) Ratio of ^18^O-labeled oxygen to ^16^O-labeled oxygen measured in cultures supplemented with ^18^O-labeled nitrate, using uninoculated medium as the control. (**C**) Concentrations of ^15^N-labeled nitrate, nitrite, and ammonium during the incubation of strains DY0037 and DY0609 in medium supplemented with ^15^N-labeled nitrate, with uninoculated medium serving as the control. (**D** and **E**) Concentrations of each inorganic nitrogen ion (NO_3_^-^, NO_2_^-^, NH_4_^+^), total inorganic nitrogen, and dissolved oxygen during cultivation of strains DY0037 (D) and DY0609 (E). ΔNH ^+^ represents the difference between the final and initial concentrations of NH ^+^ in the culture. (**F**) The proposed DNRA-dependent pathway for dark oxygen production in strains DY0037 and DY0609 involves the reduction of nitrate to nitrite by the nitrate reductase (NarGHIJK or NapADGH). An unknown enzymatic process then mediates the conversion of nitrite into oxygen and ammonium. This pathway requires two external electrons and four protons for each O_2_ molecule produced. The Gibbs free energy (ΔG) refers to the step of oxygen generation.

## Geological and ecological impacts of deep-sea microbial dark oxygen

Nodule provinces, rich in ferromanganese nodules, are well-known deep-sea geological features (*29*). Given the crucial role of oxygen in manganese nodule formation (*30*), we hypothesize that oxygen produced by *Deferribacterota* strains could potentially promote the formation of manganese nodules. Indeed, imaging revealed that *Deferribacterota* strains DY0037 and DY0609 promoted the formation of mineral precipitates in the presence of 20 mM Mn(II) (Fig. 3A and fig. S7, A and B). EDS analysis indicated that oxygen and manganese were the two most abundant elements in the precipitates formed by strains DY0037 (Fig. 3B) and DY0609 (fig. S7C). XPS analysis revealed Mn(II):Mn(IV) ratios of 46.6%:53.4% and 48.4%:51.6% in the surface of minerals formed by strains DY0037 (Fig. 3C) and DY0609 (fig. S7D), respectively. These results suggest that the oxygen produced by both strains drives the oxidation of Mn(II) to Mn(IV). EDS and XPS results consistently demonstrated the presence of MnO_2_ in the minerals, as Mn(IV) predominantly occurs in the form of MnO_2_. Since MnO_2_ is a major component of polymetallic nodules, these findings suggest a potential role for oxygen-producing microorganisms in the formation of polymetallic nodules. Building on recent reports of oxygen production within deep-sea polymetallic nodules (*19*) and our current findings, we hypothesize that oxygen-producing microorganisms may inhabit these structures, playing a role in both the formation of polymetallic nodules and the production of molecular oxygen.

**fig. S7.**
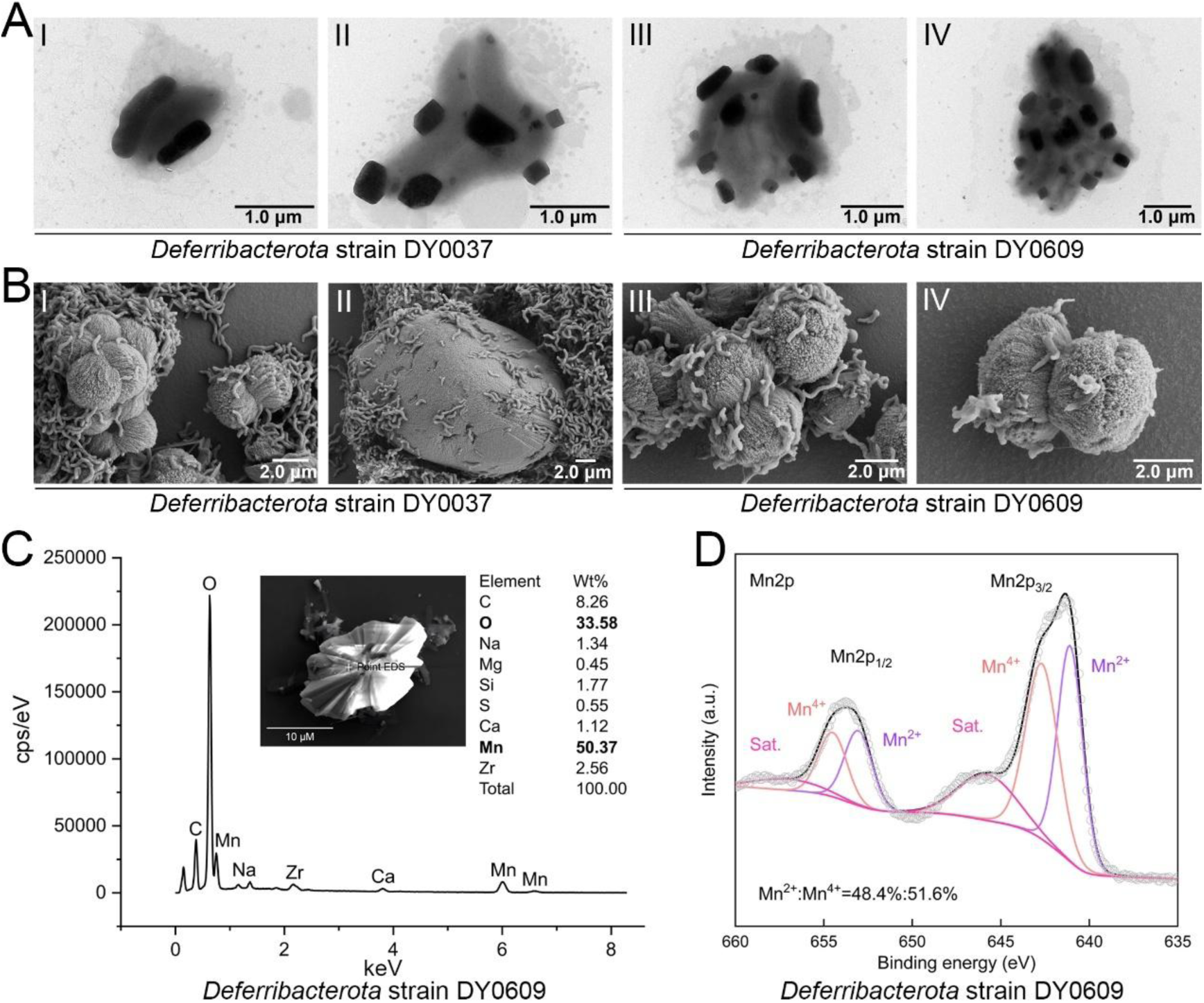
Microscopic and elemental composition analyses of minerals produced by *Deferribacterota* strains DY0037 and DY0609. (**A**) Transmission electron microscopy (TEM) images of strains DY0037 and DY0609 cultured in T16 medium with 20 mM MnCl_2_ supplementation. (**B**) Field-emission scanning electron microscopy (FESEM) images of precipitates and cell morphology of strains DY0037 and DY0609 cultured in T16 medium with 20 mM MnCl_2_ supplementation. (**C**) Energy dispersive spectrum (EDS) analysis of the precipitates formed by strain DY0609. (**D**) X-ray photoelectron spectroscopy (XPS) analysis of the ratio of manganese (II) to manganese (IV) ions in precipitate formed by strain DY0609.

**Fig. 3.**
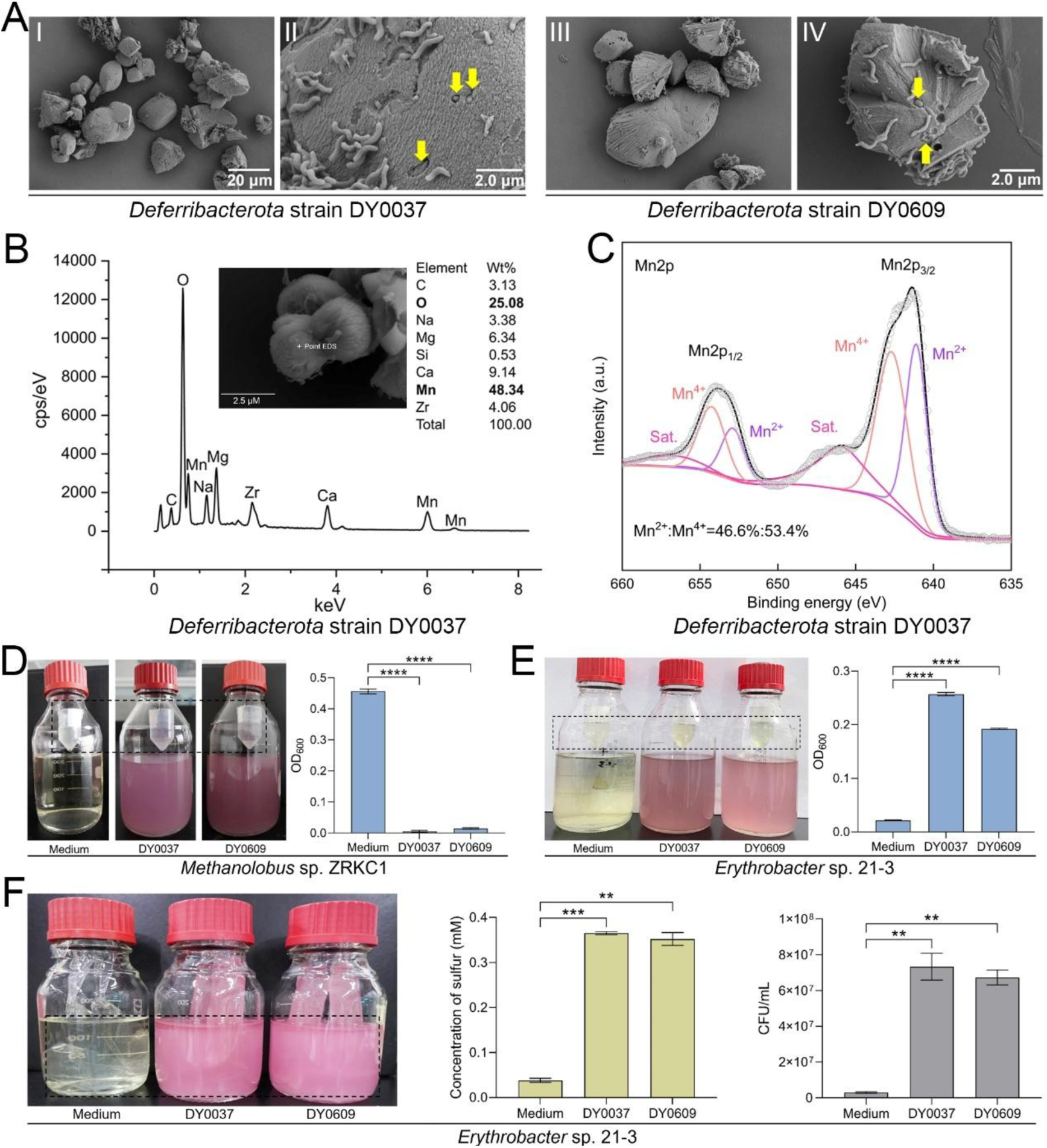
*Deferribacterota* strains potentially promotes the formation of manganese nodules and influence the growth and metabolism of other deep-sea obligately anaerobic and aerobic microbes through the oxygen they produce. (**A**) Field-emission scanning electron microscopy (FESEM) images show mineral precipitates formed by strains DY0037 and DY0609 cultured in T16 medium with 20 mM MnCl_2_ supplementation. Yellow arrows highlight cells embedded within precipitates. (**B**) Energy dispersive spectrum (EDS) analysis of the precipitates formed by strain DY0037. (**C**) X-ray photoelectron spectroscopy (XPS) analysis of the ratio of manganese (II) to manganese (IV) ion in precipitates formed by strain DY0037. (**D**) An experimental setup in which strains DY0037 and DY0609 are cultured in an anaerobic bottle, with a deep-sea obligately anaerobic methanogen (*Methanolobus* sp. ZRKC1) inoculated in a small container suspended above, to evaluate the effect of upper-layer oxygen produced by strains DY0037 and DY0609 on the growth of the anaerobe. (**E**) An experimental setup in which strains DY0037 and DY0609 are cultured in an anaerobic bottle, with a deep-sea obligately aerobic *Erythrobacter* sp. 21-3 inoculated in a small container suspended above, to evaluate the effect of upper-layer oxygen produced by strains DY0037 and DY0609 on the growth of aerobe. (**F**) An experimental setup in which strains DY0037 and DY0609 are cultured inside separate dialysis bags within an anaerobic bottle, with obligately aerobic *Erythrobacter* sp. 21-3 inoculated outside the dialysis bags using 2216E medium supplemented with 40 mM sodium thiosulfate. This setup is designed to assess the effect of the upper-layer oxygen produced by strains DY0037 and DY0609 on elemental sulfur production and the growth of *Erythrobacter* sp. 21-3.21-3. Error bars show mean ± SD. \*\**P* < 0.01, \*\*\**P* < 0.001, \*\*\*\**P* < 0.0001.

Given that strains DY0037 and DY0609 release free oxygen into the external environment, we next investigated the effect of their DOP on other deep-sea microorganisms using a two-compartment system. In this setup, an open vessel suspended above an anaerobic culture facilitated oxygen diffusion into the upper compartment (fig. S8A). The system’s functionality was confirmed by the color change of the medium in the upper vessel from colorless to pink when strains DY0037 and DY0609 were cultured in the lower compartment, indicating successful oxygen diffusion (fig. S8, B and C). Subsequent experiments demonstrated that oxygen produced by strains DY0037 and DY0609 significantly inhibited the growth of the deep-sea strictly anaerobic methanogen *Methanolobus* sp. ZRKC1 (isolated from a deep-sea cold seep) (Fig. 3D). This suggests that these strains may play a role in reducing methane production, thereby influencing methane oxidation metabolism in deep-sea environments. On the other hand, oxygen produced by strains DY0037 and DY0609 rescued the growth of a deep-sea strictly aerobic bacterium *Erythrobacter* strain 21-3 (isolated from a deep-sea cold seep) (*31*), which was cultured under strictly anaerobic conditions (Fig. 3E). When strain DY0037 or DY0609 was cultivated inside the dialysis bag and *Erythrobacter* sp. 21-3 was cultured externally, the oxygen produced by both strains significantly enhanced the production of zero-valent sulfur (ZVS), an oxidation product of thiosulfate (*31*) (Fig. 3F). This suggests that microbial DOP may play a crucial role in the deep-sea biogeochemical element cycle, particularly in the element oxidation processes.

**fig. S8.**
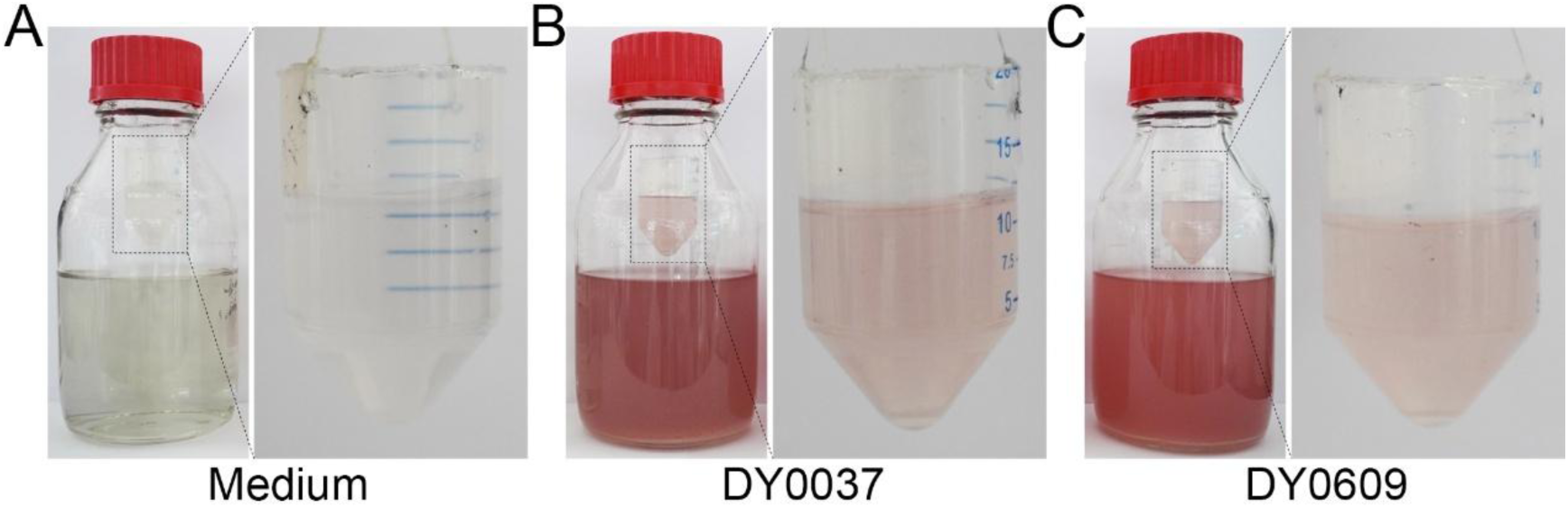
An experimental setup featuring a small container suspended inside an anaerobic bottle to confirm the presence of oxygen in the upper layer during cultivation, either without (A), or with *Deferribacterota* strain DY0037 (B), or DY0609 (C).

To mimic the impact of oxygen produced by strains DY0037 and DY0609 on the surrounding microbial community under natural conditions, we cultivated the two strains inside separate dialysis bags and simultaneously inoculated the exterior with homogenized deep-sea sediment samples (fig. S9A). Direct plating of the enriched samples onto aerobic medium revealed a substantially higher abundance of aerobic bacteria in the presence of strains DY0037 and DY0609 compared to the control group (with blank medium inside the dialysis bag) (fig. S9B). Furthermore, amplicon sequencing analysis revealed that strains DY0037 and DY0609 enriched for a diverse range of aerobic bacteria primarily (*Pseudomonadota*, including genera such as *Pseudomonas*, *Vibrio*, *Shewanella*, etc.) at significantly higher levels than the control group. In contrast, control samples predominantly enriched anaerobic bacteria, mainly members of *Clostridia* and *Fusobacteriota* (fig. S9C). The analysis of aerobic metabolism-associated genes (*32*) further revealed a significantly higher abundance in the experimental groups (strains DY0037 and DY0609) compared to the controls (fig. S9D). These results suggest that oxygen produced by *Deferribacterota* strains may create an oxic niche in deep-sea environments, facilitating the persistence of aerobic life. This observation also offers a mechanistic explanation for the seemingly paradoxical presence of aerobic microbes in nominally anoxic environments (such as groundwater, subsurface, lake, wetland, deep sea, etc.) (*4, 14*), and suggests that similar microbial DOP processes may be widespread in these environments.

**fig. S9.**
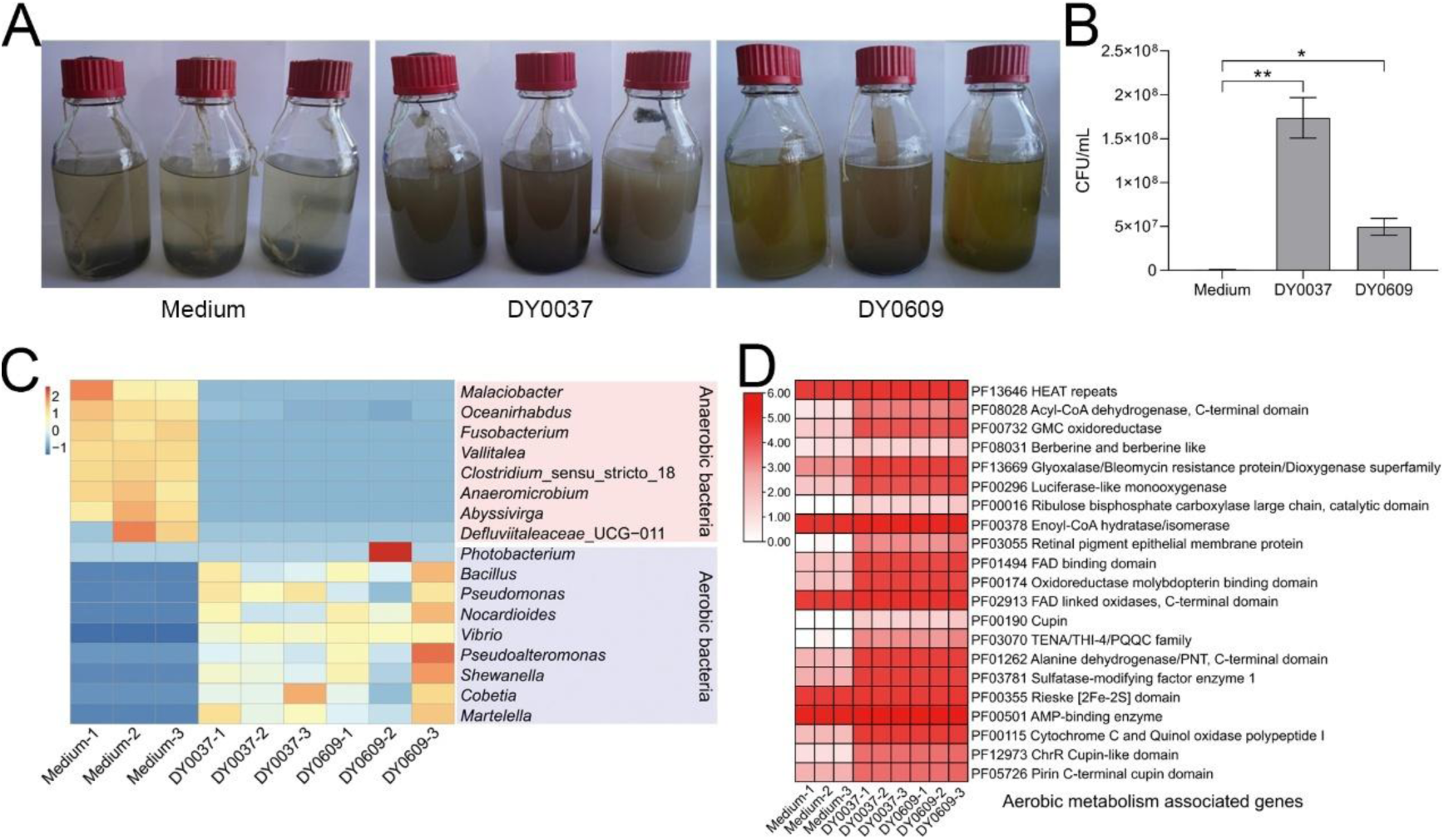
Effects of *Deferribacterota* strains DY0037 and DY0609 on the microbial community composition and aerobic metabolism in deep-sea sediment. (**A**) An experimental setup in which strains DY0037 and DY0609 are cultured within a dialysis bag inside an anaerobic bottle, with an equal amount of sediment sample inoculated outside the dialysis bag, and uninoculated medium serving as the control. (**B**) Quantification of aerobic colony-forming units (CFUs) in experimental groups (strains DY0037 and DY0609) and the control group (uninoculated medium) using the plate count method. (**C**) Amplicon sequencing analysis of the enriched microbial community composition (aerobes and anaerobes) in experimental groups (strains DY0037 and DY0609) and the control group (uninoculated medium). (**D**) Abundance of genes associated with aerobic metabolism in the enriched microbial communities from experimental groups (strains DY0037 and DY0609) and the control group (uninoculated medium), assessed by amplicon sequencing. Error bars show mean ± SD. \**P* < 0.05, \*\**P* < 0.01.

In addition to supporting aerobic microbial life, what is the potential contribution of *Deferribacterota* DOP to animal respiration? To assess the role of oxygen produced by strains DY0037 and DY0609 in supporting animal survival, we used ants (fig. S10A) and locusts (fig. S10C) as experimental organisms. We observed a significant extension in both ant and locust survival times in the presence of oxygen diffused from cultures of strains DY0037 and DY0609, with strain DY0609 exhibiting a more pronounced effect (fig. S10, B and D). This result is consistent with its relatively higher oxygen production capability throughout the entire cultivation period (Fig. 1C). These findings demonstrate that oxygen production by strains DY0037 and DY0609 could have broad implications for both microbial and macroscopic life, suggesting potential applications in life support systems for space and deep-sea exploration. Notably, *Deferribacterota* bacteria have been shown to dominate the gut microbiome of blind shrimp inhabiting hydrothermal vents (*25*). The symbiotic relationship between *Deferribacterota* bacteria and animal host, evidenced by the exchange of material and energy, suggests that *Deferribacterota* bacteria function as gut symbionts (*25*). The potential for *in situ* oxygen production by this symbiont in blind shrimp offers a significant advantage for their survival in extreme environments and provides new insights into the interaction between gut symbionts and their host.

**fig. S10.**
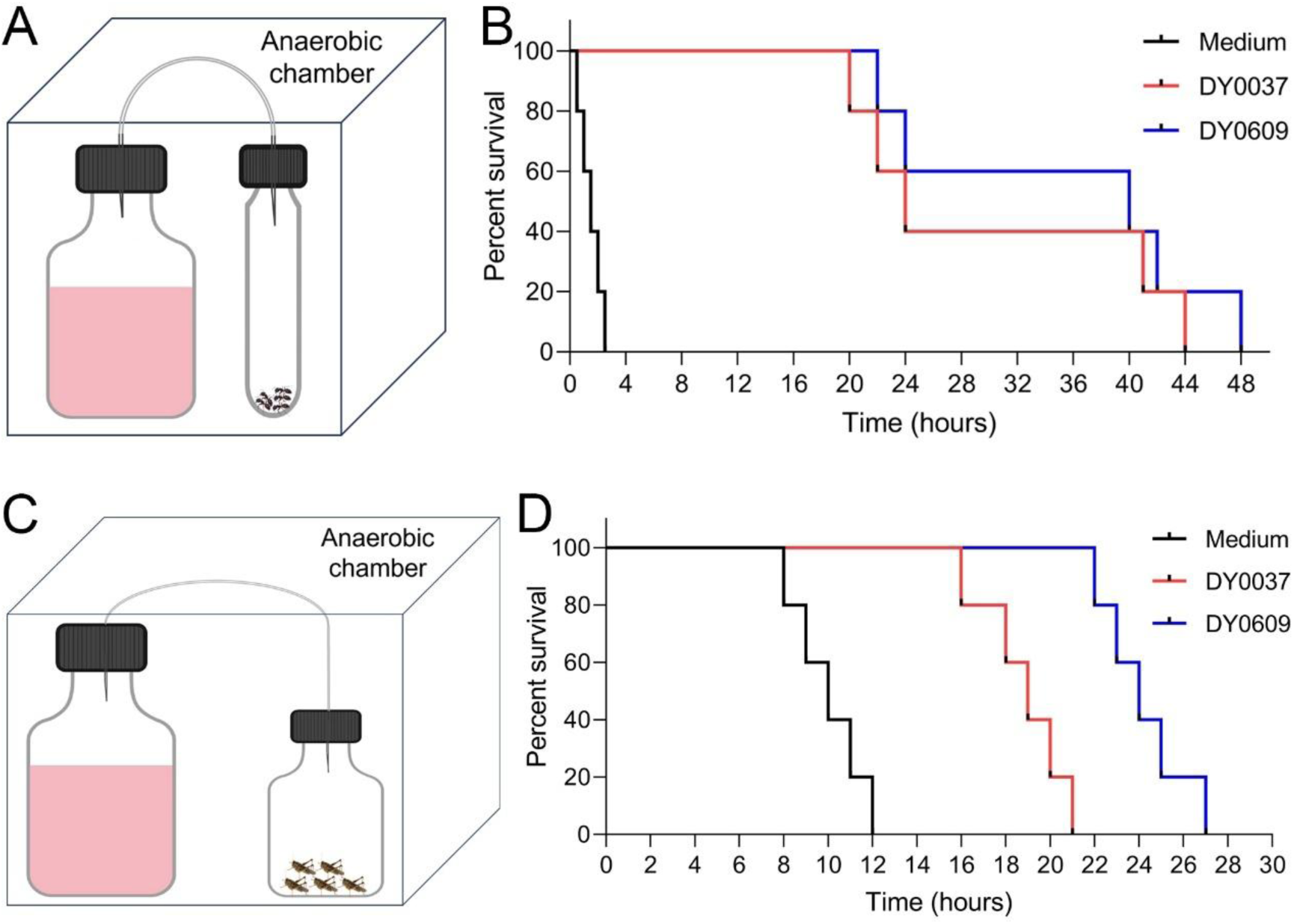
The supporting effect of oxygen produced by *Deferribacterota* strains DY0037 and DY0609 on insect survival when cultured in an oxygen-deprived container. (**A**) An experimental setup was designed to assess the impact of oxygen produced by strains DY0037 and DY0609 on ant survival time. It consisted of a 500 mL anaerobic bottle connected to a 15 mL anaerobic tube within an anaerobic chamber. (**B**) Survival time assays of ants exposed to oxygen produced by strains DY0037 and DY0609. (**C**) An experimental setup was designed to assess the impact of oxygen produced by strains DY0037 and DY0609 on locust survival time. It consisted of a 500 mL anaerobic bottle connected to a 250 mL anaerobic bottle within an anaerobic chamber. (**D**) Survival time assays of locusts exposed to oxygen produced by strains DY0037 and DY0609.

## Diverse deep-sea microorganisms produce dark oxygen

To investigate the prevalence of deep-sea microbial DOP, we performed targeted enrichments from hydrothermal vent, cold seep, and seamount samples using a nitrate-containing medium. This led to the isolation of a diverse collection of oxygen-producing microbes. These included some *Pseudomonadota* strains DY0742 (from hydrothermal vent), 24LQ007 (from cold seep), and 273 (from seamount), *Campylobacterota* strain DY0563 (from hydrothermal vent), and *Deinococcota* strain DY0809b (from hydrothermal vent). Genomic and physiological assays ruled out the possibility of photosynthetic and other abiotic oxygen production in these strains, confirming their DOP capability. Further genomic analysis of the *Pseudomonadota* strains (DY0742, 24LQ007, and 273) revealed the presence of complete DNRA and denitrification pathways (Fig. 4A). During anaerobic growth of these *Pseudomonadota* strains, nitrate concentrations decreased significantly, while nitrite concentrations increased, with no notable change in ammonium levels. Additionally, a reduction in total inorganic nitrogen content was observed, suggesting nitrogen loss as gaseous products during denitrification (Fig. 4B and fig. S11). Additionally, dissolved oxygen concentrations in cultures of these three *Pseudomonadota* strains reached approximately 20 μM during growth. Previous studies have reported oxygen production via denitrification in *Pseudomonas* species (*20*). Based on our findings, we propose that these deep-sea *Pseudomonadota* strains are capable of oxygen production through denitrification. To validate the DOP pathway, we generated a *norBCD* deletion mutant of *Pseudomonadota* strain DY0742, as this gene cluster is potentially associated with oxygen production. Under anaerobic conditions, the wild-type strain DY0742 exhibited robust growth, while the mutant failed to grow. However, both strains showed normal growth under aerobic conditions (Fig. 4C). This suggests that *norBCD* is essential for anaerobic denitrification and oxygen production in strain DY0742, with its deletion impairing anaerobic growth.

**fig. S11.**
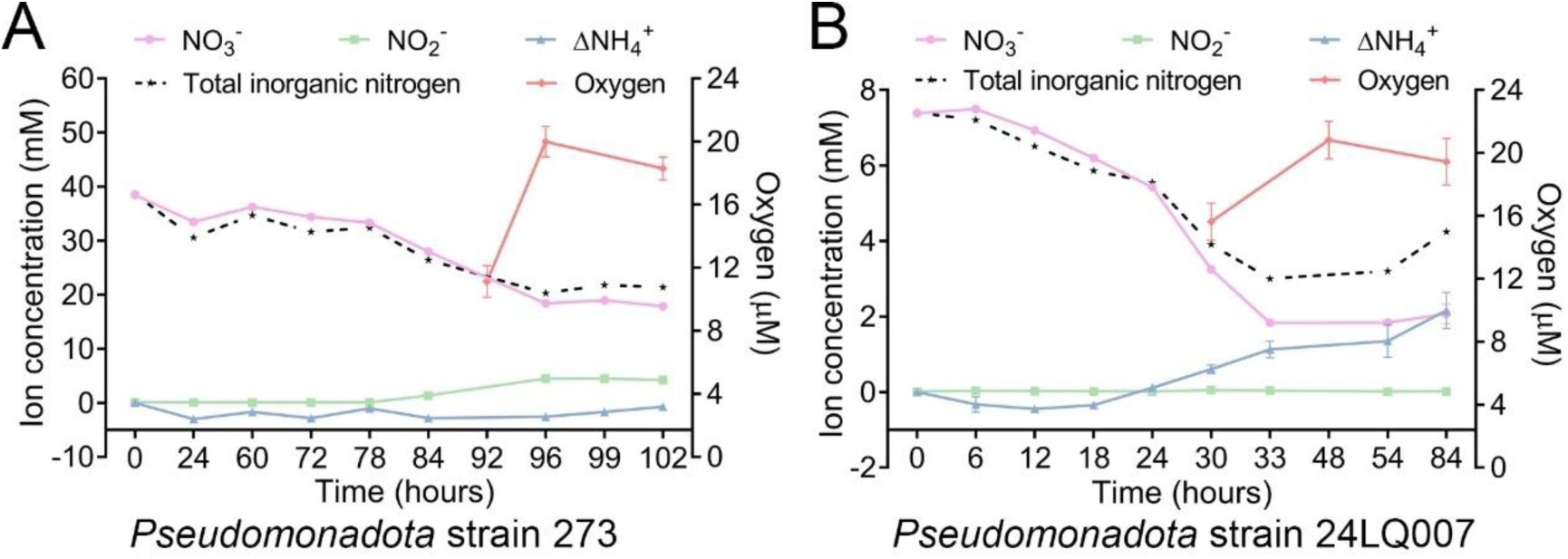
Dynamics of concentrations of each inorganic nitrogen ion (NO ^-^, NO ^-^, or NH ^+^), total inorganic nitrogen, and dissolved oxygen during cultivation of *Pseudomonadota* strains 273 (A) and 24LQ007 (B). ΔNH ^+^ indicates the difference between final and initial concentrations of NH ^+^ present in the culture.

**Fig. 4.**
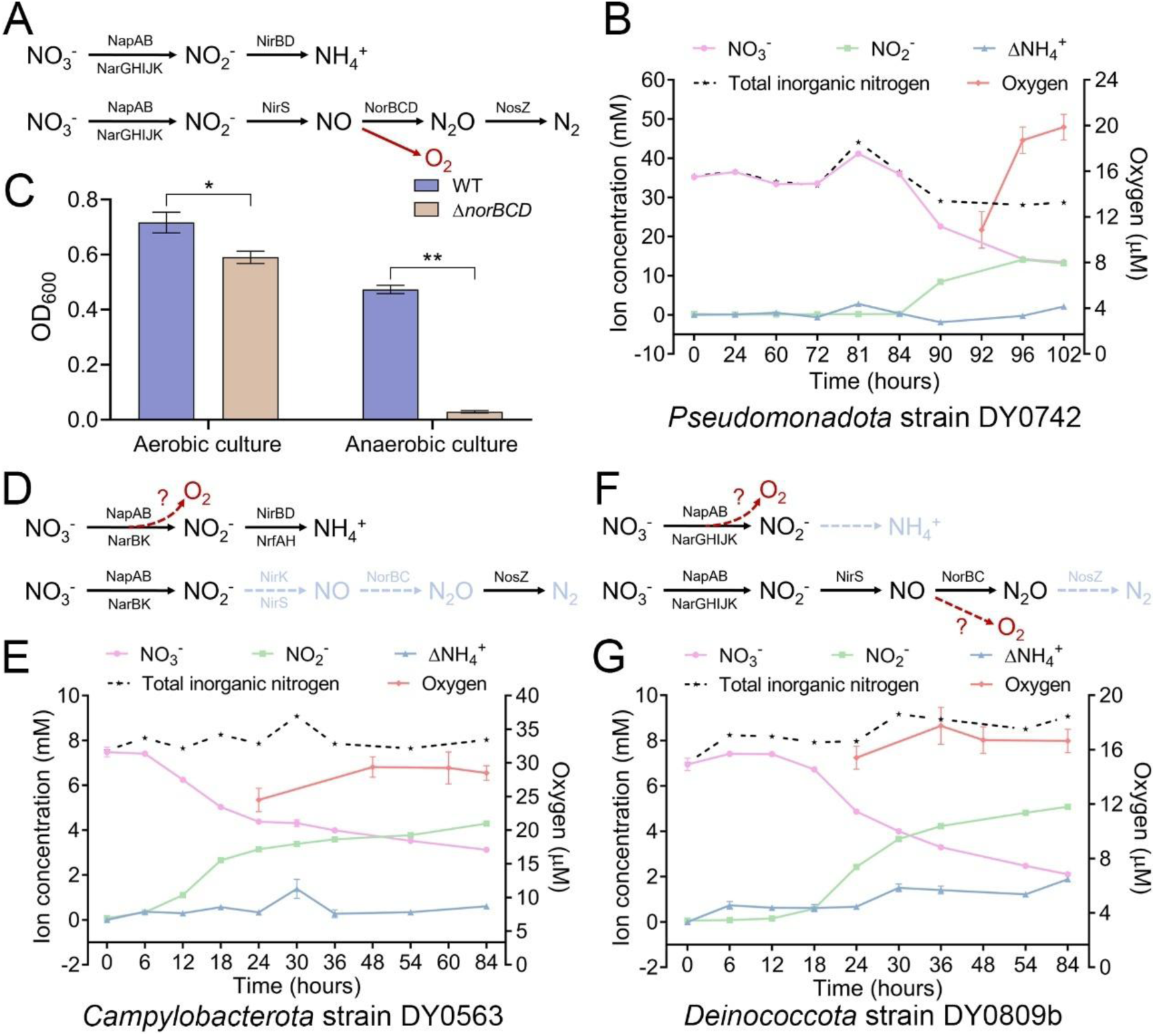
Genomic and metabolic profiles of diverse deep-sea bacteria that produce dark oxygen through nitrate reduction-dependent pathways. (**A, D, F**) Analysis of genes involved in the DNRA and denitrification pathways, along with the proposed oxygen production pathway, in *Pseudomonadota* strain DY0742 (A), *Campylobacterota* strain DY0563 (D), and *Deinococcota* strain DY0809b (F). Solid black arrows indicate the presence of relevant genes and intermediates, while dashed light blue arrows represent the absence of these genes and intermediates. Solid or dashed red arrows indicate the potential locations of oxygen production. (**B, E, G**) Concentrations of each inorganic nitrogen ion (NO_3_ ^-^, NO_2_ ^-^, NH_4_ ^+^), total inorganic nitrogen, and dissolved oxygen during cultivation of *Pseudomonadota* strain DY0742 (B), *Campylobacterota* strain DY0563 (E), and *Deinococcota* strain DY0809b (G). ΔNH_4_ ^+^ indicates the difference between the final and initial concentrations of NH_4_ ^+^ present in the culture. (**C**) Growth profile of wild-type strain DY0742 and its *norBCD* deletion mutant under aerobic and anaerobic conditions. Error bars show mean ± SD. \**P* < 0.05, \*\**P* < 0.01.

Genomic analyses revealed that the *Campylobacterota* strain DY0563 possesses a complete DNRA pathway and an incomplete denitrification pathway (Fig. 4D), while the *Deinococcota* strain DY0809b contains an incomplete DNRA pathway and a nearly complete denitrification pathway, lacking only the *nosZ* gene (Fig. 4F). Ion concentration analyses revealed a significant decrease in nitrate concentrations, accompanied by increases in nitrite and dissolved oxygen concentrations, during the growth of both *Campylobacterota* strain DY0563 and *Deinococcota* strain DY0809b. No substantial change in ammonium concentrations was observed (Fig. 4, E-G). Therefore, we propose that *Campylobacterota* strain DY0563 and *Deinococcota* strain DY0809b may generate oxygen during the reduction of nitrate to nitrite (Fig. 4D), with the *Deinococcota* strain DY0809b potentially producing oxygen through a denitrification pathway similar to that of *Pseudomonadota* bacteria (Fig. 4F). However, the exact mechanism of oxygen production in these strains remains unclear, necessitating further investigation to elucidate the underlying processes and identify the key genes involved.

Furthermore, in addition to the aforementioned oxygen-producing microorganisms, we conducted a comparative genomic analysis of nitrogen metabolism genes in other deep-sea microbial groups that do not exhibit nitrate-dependent oxygen production. Our analysis revealed that microorganisms capable of producing oxygen predominantly harbor genes associated with either the DNRA or denitrification pathways, whereas those unable to produce oxygen through nitrate reduction generally lack these genes (fig. S12). Given the widespread presence of the DNRA and denitrification pathways in many microorganisms, we infer that oxygen production driven by nitrate reduction may be broadly distributed across diverse deep-sea environments, contributing to the availability of oxygen in surrounding habitats.

**fig. S12.**
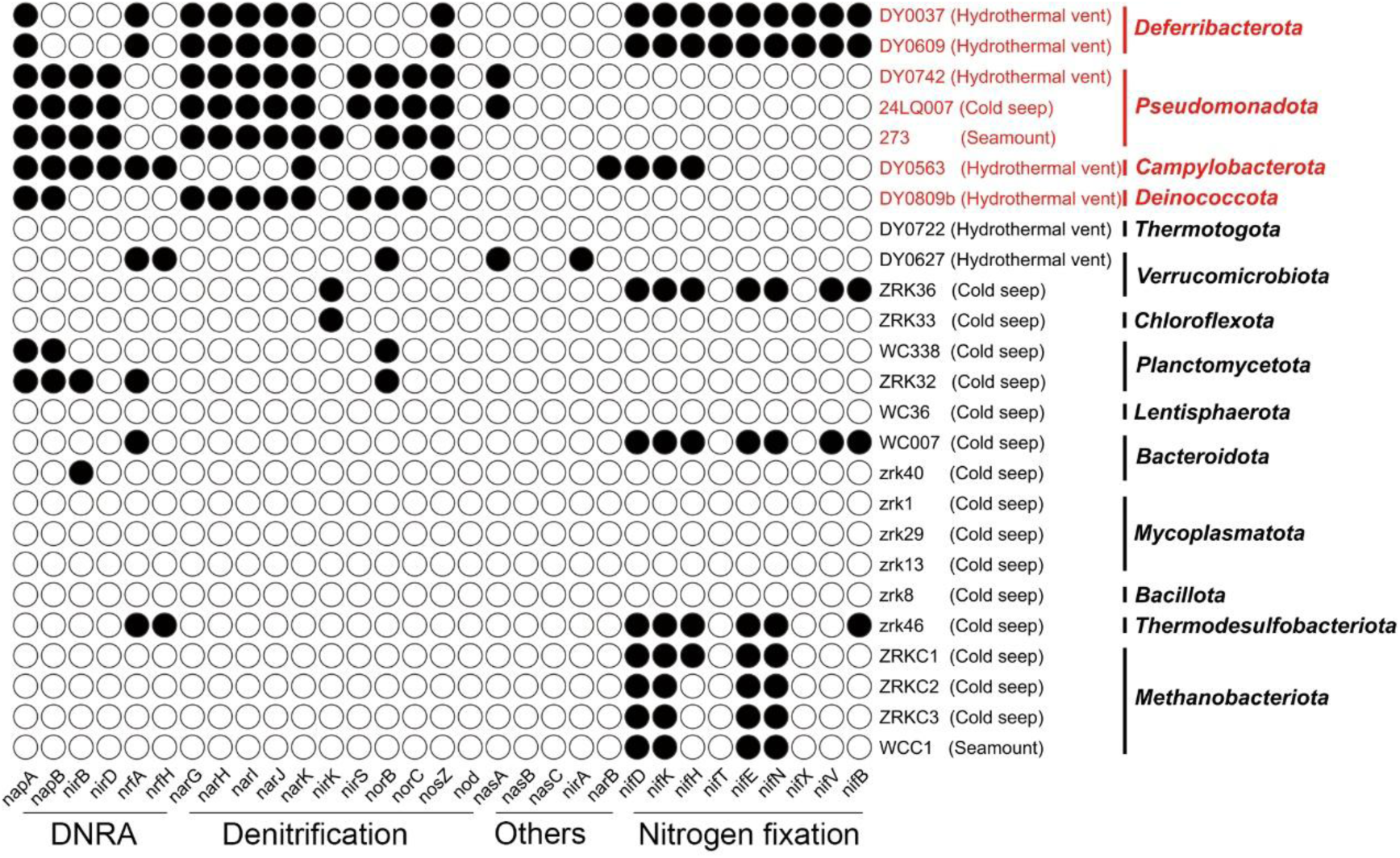
Distribution of nitrogen metabolism-associated genes across some deep-sea microbial representatives. Statistical analysis of the distribution of genes associated with nitrogen metabolism—including DNRA (dissimilatory nitrate reduction to ammonium), denitrification, nitrogen fixation, and others—across cultured deep-sea microbial representatives in our laboratory. Microbial groups capable of producing dark oxygen through nitrate reduction pathways are highlighted in red, while those lacking this capability are shown in black. The isolation source of each strain is indicated in parentheses, and the corresponding bacterial phylum is listed after the vertical line.

## Conclusions and perspectives

This study presents the cultivation of diverse deep-sea microbial groups capable of producing dark oxygen through various nitrate reduction pathways, including the identification of a novel oxygen-producing pathway coupled to DNRA in the *Deferribacterota* phylum. Our findings strongly suggest that microbial oxygen production in the deep sea is closely linked to nitrate metabolism. Consistently, oxygen concentrations with depth in the deep sea (>1000 m) were positively correlated with available nitrate levels (*21*). Thus, microbial dark oxygen could serve as a significant oxygen source in nitrate-rich ocean worlds, even in the absence of light. This oxygen supports the growth of the faunal community, influences the growth and metabolism of both aerobic and anaerobic microorganisms, and shapes the geological system by promoting the formation of polymetallic nodules (Fig. 5A). Given the dark oxygen-producing capability of polymetallic nodules (*19*), mining these resources could potentially disrupt deep-sea ecosystems, thereby limiting the exploitation of these valuable mineral deposits. Microorganisms that produce dark oxygen, as cultured in our study, could serve as promising candidates for remediating deep-sea ecological environments damaged by polymetallic nodule mining. Furthermore, these dark oxygen-producing microorganisms have the potential to enhance oxygen levels in ocean oxygen minimum zones.

**Fig. 5.**
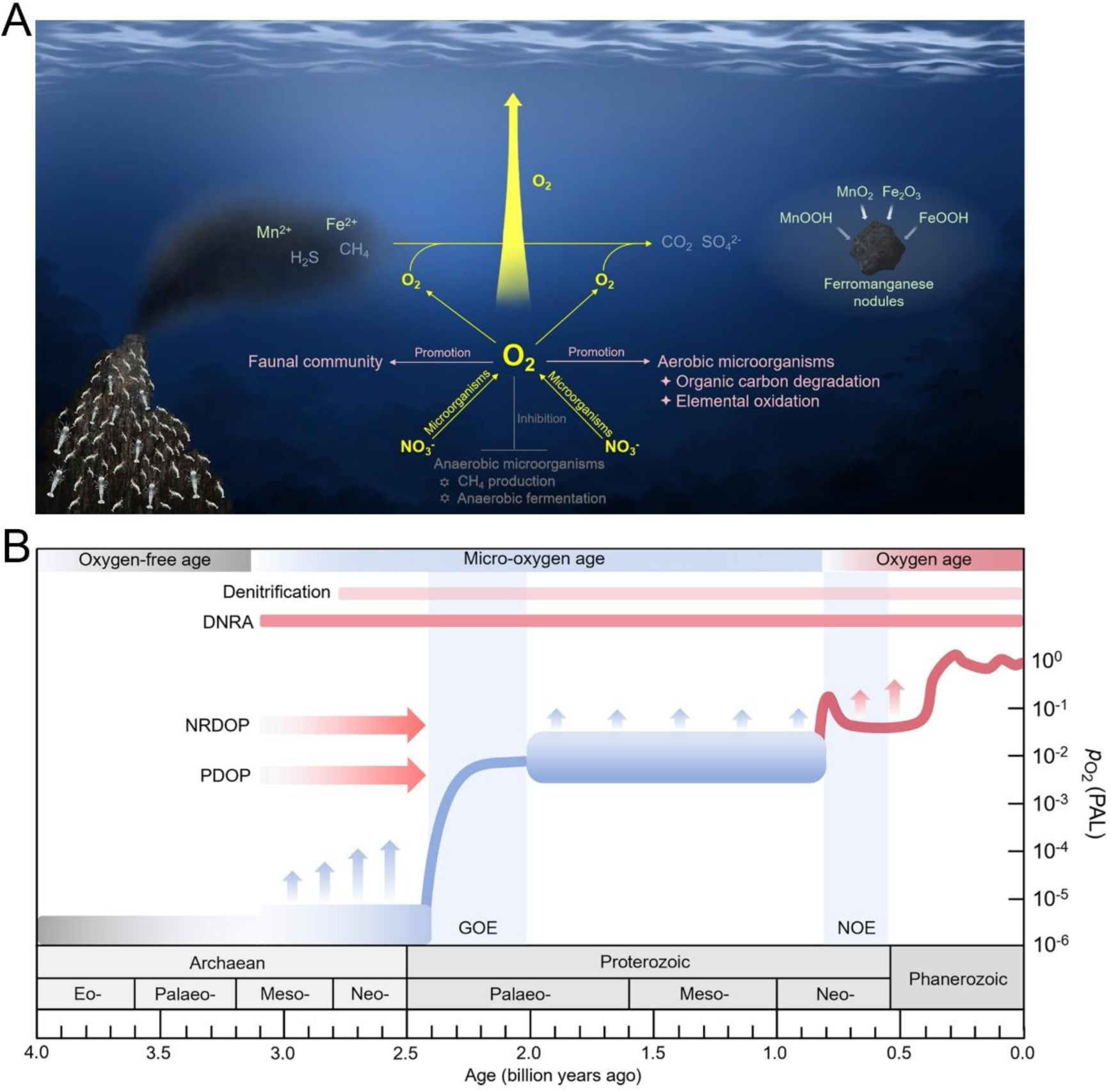
The proposed models illustrate the contribution of microbial dark oxygen produced through nitrate reduction pathways to both deep-sea ecosystems and Earth’s atmospheric oxygen levels. **(A**) A schematic diagram depicts the ecological impact of dark oxygen generated by deep-sea microorganisms through nitrate reduction pathways. Yellow arrows represent the release of oxygen by these microorganisms during nitrate reduction. (**B**) A proposed model outlines the contributions of various oxygen-producing pathways to the evolution of Earth’s atmospheric oxygen (O₂) levels over geological time. The gray, blue, and red bands are based on references (*41*) and (*42*). *p*O2 (PAL) represents the atmospheric partial pressure of oxygen relative to the present atmospheric level. The Great Oxygenation Event (GOE), occurring approximately 2.5–2.4 to 2.1–2.0 billion years ago, and the Neoproterozoic Oxygenation Event (NOE), spanning 0.8 to 0.5 billion years ago, are highlighted as key intervals of atmospheric oxygen increase (see references (*43*), (*44*), and (*45*) for reviews of NOE through Devonian oxygenation). The light red and red shaded boxes in the diagram indicate the respective timeframes for the emergence of microbial denitrification and DNRA pathways. Red arrows point to the right, indicating the approximate timing of nitrate reduction-dependent oxygenic production (NRDOP) and photosynthesis-dependent oxygenic production (PDOP), though their exact ages are uncertain and not plotted relative to the x-axis.

In deep-sea environments, DNRA is more prevalent than denitrification among microbial communities (*33–35*). Our research on deep-sea hydrothermal vent *Deferribacterota* bacteria, which link DNRA to DOP, highlights their considerable contribution to early ocean oxygen sources. Moreover, previous findings suggest substantial atmospheric oxygen levels existed approximately 3 billion years ago, more than 600 million years before the GOE and 300-400 million years earlier than prior estimates of Earth’s surface oxygenation (*36*). Given the presence of nitrogen oxides on early Earth, our results, in conjunction with previous findings, offer a novel perspective on the origins of oxygen (*6, 7*). Intra-aerobic denitrification and anaerobic DNRA may have either preceded photosynthetic oxygen production or created critical niches for the evolution of aerobic pathways in the largely anaerobic environment before the GOE. Evidence from Archean and early Proterozoic geochemical records also suggests that DNRA, rather than solely denitrification to N_2_, may have contributed to biological nitrate utilization (37) (Fig. 5B). Therefore, both nitrate reduction-dependent oxygenic production (NRDOP) and photosynthesis-dependent oxygenic production (PDOP) are proposed to play critical roles in the rise of the GOE.

Furthermore, both nitrate and chlorate have been detected on Mars and in Martian meteorites, with nitrate levels reaching up to 1000 ppm in Martian sediments (*38, 39*), suggesting potential resources for life. The subsurface nitrates on Mars could serve as a source of bioavailable nitrogen and oxygen, particularly within its subsurface ecosystems (*40*). he dark oxygen-producing bacteria, including strains from *Deferribacterota*, *Deinococcota*, *Campylobacterota*, and *Pseudomonadota*, isolated from deep-sea hydrothermal vents, are capable of anaerobic growth and can tolerate a wide range of temperatures. These traits indicate their potential to generate dark oxygen from nitrate on Mars, which could contribute to terraforming efforts and potentially support human colonization in the future.

## Materials and methods

### Enrichment and cultivation of deep-sea oxygen-producing microorganisms

The deep-sea hydrothermal vent samples were collected by ShenhaiYihao from the “lost city” hydrothermal field in the North Atlantic (W42°7’9.38”, N 30°7’25.92”). The deep-sea cold seep (E119°17’07.322”, N22°06’58.598”) and seamount (E110°41’12.616”, N17°22’11.438”) samples were collected by *RV KEXUE* in the South China Sea. To isolate and cultivate oxygen-producing microorganisms, these deep-sea sedimental samples were added to the 15 mL anaerobic tube containing 10 mL T13 medium (0.02 g/L Na_2_SO_4_, 1.0 g/L KH_2_PO_4_, 1.0 g/L NH_4_Cl, 20.0 g/L NaCl, 3.0 g/L MgCl_2_, 0.5 g/L KCl, 0.15 g/L CaCl_2_, 1.0 g/L NaNO_3_, 1.0 g/L CH_3_COONa, 1.0 g/L NaHCO_3_, 0.5 g/L fumarate, 0.5 g/L yeast extract, 0.5 g/L peptone, 1.0 g/L cysteine hydrochloride, 500 µL/L 0.1 % (w/v) resazurin, 10 mL/L trace element solution (1.5 g/L nitrilotriacetic acid, 3.0 g/L MgSO_4_**·**7H_2_O, 0.5 g/L MnSO_4_**·**H_2_O, 1.0 g/L NaCl, 0.1 g/L FeSO_4_**·**7H_2_O, 0.18 g/L CoSO_4_**·**7H_2_O, 0.1 g/L CaCl_2_**·**2H_2_O, 0.18 g/L ZnSO_4_**·**7H_2_O, 0.01 g/L CuSO_4_**·**5H_2_O, 0.02 g/L AlK(SO_4_)_2_**·**12H_2_O, 0.01 g/L H_3_BO_3_, 0.01 g/L Na_2_MoO_4_**·**2H_2_O, 0.03 g/L NiCl_2_**·**6H_2_O, 0.3 mg/L Na_2_SeO_3_**·**5H_2_O, 0.4 mg/L Na_2_WO_4_**·**2H_2_O, 1.0 L distilled water), 1.0 L sterilized distilled water, pH 7.0), respectively. The medium was prepared under a 100% N_2_ gas phase and sterilized by autoclaving at 115 °C for 30 minutes. The inoculated media were anaerobically incubated at either 28 °C or 55 °C for one week. T13 medium, supplemented with 15 g/L agar, was evenly spread onto the inner wall of a Hungate tube, creating a thin layer for bacterial growth. Following this, 50 µL of the enriched culture was anaerobically transferred to an anaerobic roll tube and spread onto the medium layer. These tubes were also anaerobically incubated at either 28 °C or 55 °C for three days. Single colonies were selected using sterilized bamboo sticks and cultured in 15 mL Hungate tubes containing 10 mL T13 medium at 28 °C or 55 °C for three days. Tubes in which the culture medium turned pink were selected for microbial identification by PCR. Deep-sea microorganisms were isolated and purified through repeated use of the Hungate roll-tube method. Purity was regularly confirmed through observation with a transmission electron microscope (TEM) and partial sequencing of the 16S rRNA gene.

### TEM observation

To observe the morphological characteristics of *Deferribacterota* strains DY0037 and DY0609, 10 mL of culture was collected by centrifugation at 5000 × *g* for three minutes. The cells were then washed three times with PBS buffer (137 mM NaCl, 2.7 mM KCl, 10 mM Na₂HPO₄, 1.8 mM KH₂PO₄, pH 7.4). After washing, the cells were resuspended in 20 μL of PBS buffer and transferred onto copper grids coated with a carbon film by immersing the grids in the cell suspension for 20 minutes. All samples were examined using a transmission electron microscope (TEM, HT7700, Hitachi, Japan).

### Genome sequencing, annotation, and analysis

For genomic sequencing, *Deferribacterota* strains DY0037 and DY0609, *Campylobacterota* strain DY0563 and *Deinococcota* strain DY0809b were incubated in QY5 medium (0.17 g/L KH_2_PO_4_, 3.0 g/L NaNO_3_, 9.0 g/L NaCl, 1.0 g/L MgCl_2_, 2.5 g/L D-Trehalose anhydrous, 0.17 g/L KCl, 0.08 g/L CaCl_2_, 0.17 g/L NH_4_Cl, 1.0 g/L NaHCO_3_, 0.5 g/L yeast extract, 2.5 g/L peptone, 0.2 g/L cysteine hydrochloride, 500 µL/L 0.1 % (w/v) resazurin, 5 mL/L trace element solution, 0.5 L sterilized distilled water and 0.5 L sterilized seawater, pH 7.0); *Pseudomonadota* strains DY0742, 24LQ007, and 273 were incubated in T14 medium (0.02 g/L Na_2_SO_4_, 1.0 g/L KH_2_PO_4_, 1.0 g/L NH_4_Cl, 20.0 g/L NaCl, 3.0 g/L MgCl_2_, 0.5 g/L KCl, 0.15 g/L CaCl_2_, 1.0 g/L NaNO_3_, 1.0 g/L CH_3_COONa, 1.0 g/L NaHCO_3_, 0.5 g/L fumarate, 0.5 g/L yeast extract, 0.5 g/L peptone, 1.0 g/L cysteine hydrochloride, 500 µL/L 0.1 % (w/v) resazurin, 10 mL/L trace element solution, 1.0 L sterilized distilled water, pH 7.0). Bacterial cells were harvested after three days of incubation at either 28 °C (strains DY0037, DY0609, DY0742, DY0828, LQ001, and 273), 37 °C (strain DY0563), or 55 °C (strain DY0809b). Genomic DNA was isolated by using the PowerSoil DNA isolation kit (Mo Bio Laboratories Inc., Carlsbad, CA). Thereafter, the genome sequencing was carried out with both the Illumina NovaSeq PE150 (San Diego, USA) and Nanopore PromethION platform (Oxford, UK) at the Beijing Novogene Bioinformatics Technology Co., Ltd. A complete description of the library construction, sequencing, and assembly was performed as previously described (*46*). We used seven databases to predict gene functions, including Pfam (Protein Families Database, http://pfam.xfam.org/), GO (Gene Ontology, http://geneontology.org/) (*47*), KEGG (Kyoto Encyclopedia of Genes and Genomes, http://www.genome.jp/kegg/) (*48*), COG (Clusters of Orthologous Groups, http://www.ncbi.nlm.nih.gov/COG/) (*49*), NR (Non-Redundant Protein Database databases), TCDB (Transporter Classification Database), and Swiss-Prot (http://www.ebi.ac.uk/uniprot/) (*50*). A whole genome Blast search (E-value less than 1e-5, minimal alignment length percentage larger than 40%) was performed against above seven databases.

### Phylogenetic analysis

To construct a maximum likelihood 16S rRNA phylogenetic tree, the full-length 16S rRNA gene sequences of *Deferribacterota* strains DY0037 and DY0609 and other related taxa were extracted from their corresponding genomes (www.ncbi.nlm.nih.gov/). The phylogenetic tree was constructed by the W-IQ-TREE web server (http://iqtree.cibiv.univie.ac.at) (*51*) using the “GTR+F+I+G4” model, and the Interactive Tree of Life (iTOL v5) online tool (*52*) was used to edit the phylogenetic trees.

### Measurement of bacterial growth and dissolved oxygen concentration

To assess the growth and dissolved oxygen production of *Deferribacterota* strains DY0037 and DY0609, T13 medium was used. For *Campylobacterota* strain DY0563 and *Deinococcota* strain DY0809b, QY5 medium was employed, while T14 medium was used for *Pseudomonadota* strains 24LQ007, DY0742, and 273. For each assay, 4.0 mL of fresh culture from different strains was inoculated into a 500 mL Hungate bottle containing 400 mL of the respective medium. The Hungate bottles were anaerobically incubated at 28 °C (for strains DY0037, DY0609, DY0742, DY0828, LQ001, and 273), 37 °C (for strain DY0563), or 55 °C (for strain DY0809b). Bacterial growth was monitored daily by measuring OD_600_ values using a microplate reader (Infinite M1000 Pro; Tecan, Männedorf, Switzerland) until the cultures reached stationary phase. Additionally, to investigate the effect of nitrate on the growth of strains DY0037 and DY0609, their growth in T13 medium with and without sodium nitrate was compared. Three replicates were performed for each condition. The dissolved oxygen concentrations were measured using a liquid-phase oxygen electrode (Hansatech, Oxygraph+, UK).

### Experimental exclusion of alternative oxygen sources

To investigate the effect of seed culture components on oxygen production in *Deferribacterota* strains DY0037 and DY0609, seed culture was incorporated into T13 medium both before and after sterilization. After two days of incubation at 28 °C, growth and dissolved oxygen levels were measured. To rule out H₂O₂ dismutation as a potential mechanism for oxygen production, strains DY0037 and DY0609 were cultured in T13 medium supplemented with 0.2 mM pyruvate, as α-keto acids can detoxify H₂O₂ through abiotic decarboxylation (*24*). Additionally, to exclude photosynthetic oxygen production, the strains were grown under both light and dark conditions at 28 °C for two days, with growth and dissolved oxygen levels measured thereafter.

### Transcriptomics

For transcriptomic sequencing, *Deferribacterota* strains DY0037 and DY0609 were cultured in 400 mL of T13 medium supplemented with varying concentrations of sodium nitrate (0.1 g/L, 0.5 g/L, 1.0 g/L, and 3.0 g/L) at 28 °C for two days. Three biological replicates were cultured for each condition. Afterward, cells from each replicate were collected and sent for transcriptomic sequencing to Novogene (Tianjin, China), as previously described (*46, 53*). Briefly, RNA-seq data were generated using a NovaSeq 6000 instrument with PE150. Raw reads were trimmed for adaptors and low-quality sequences using Trimmomatic (*54*), and mRNA was recovered with SortMeRNA (*55*) (default settings) after removing tRNA and rRNA. The activity of strains DY0037 and DY0609 was evaluated using their complete circular genomes, and gene transcription levels were quantified using Burrows-Wheeler Aligner (BWA, v. 0.7.17-r1188) (*56*) to map reads, followed by SAMtools (v. 1.13) (*56*) conversion to BAM files. Read coverage was calculated with BEDTools (v. 2.30.0) (*57*), normalized to genome length, and gene expression was quantified using Fragments Per Kilobase Million (FPKM). The transcribed rank of genes was then calculated based on log_2_[FPKM] values.

### Verification of oxygen production from nitrate using a stable isotope assay

To investigate whether the oxygen originated from nitrate, *Deferribacterota* strains DY0037 and DY0609 were cultured in T13 medium containing 10 mM ^18^O-labeled KNO_3_ instead of 1.0 g/L NaNO_3_. After three days of incubation, the headspace gas was collected and analyzed for ¹⁸O₂ and ¹⁶O₂ using an isotope mass spectrometer (Delta Q, Thermo Fisher, USA).

### Ion concentration detection

To quantify the ion concentrations (NO₃⁻, NO₂⁻, and NH₄⁺) during the growth of *Deferribacterota* strains DY0037 and DY0609 in T13 medium, culture supernatants were collected by centrifugation at 12,000 × *g* for 10 minutes every 12 hours over a four-day period. For the other bacterial strains (*Campylobacterota* strain DY0563, *Deinococcota* strain DY0809b, and *Pseudomonadota* strain 24LQ007 in QY5 medium, and *Pseudomonadota* strains DY0742 and 273 in T14 medium), culture supernatants were collected after 84 or 102 hours of incubation. These supernatants were then analyzed for ion concentrations (NO₃⁻, NO₂⁻, and NH₄⁺) using a continuous flow analyzer (QuAAtro39, Seal, Germany).

### Verification of NO_2_^-^/NH_4_^+^ production from nitrate using a stable isotope assay

To determine whether nitrate can produce ammonium through the DNRA pathway, *Deferribacterota* strains DY0037 and DY0609 were cultured in T13 medium containing 10 mM ¹⁵N-labeled KNO₃ instead of 1.0 g/L NaNO₃. After three days of incubation, the culture supernatants were collected and analyzed using a Gas Isotope Ratio Mass Spectrometer (MAT 253, Thermo Fisher, USA) to detect and quantify ammonium.

### X-ray diffraction (XRD) assay

To determine the mineral composition produced by *Deferribacterota* strains DY0037 and DY0609 during their growth phase, 2 g of mineral particles were collected from each strain. These particles were washed three times with distilled water and then thoroughly dried. XRD analysis of the dried mineral powders was performed using a Rigaku SmartLab diffractometer (SmartLab 9KW, Rigaku Corporation, Japan) under the following conditions: 9 kW, 45 kV, 200 mA, with a step size of 0.02°, a scanning speed of 20°/min, and a 2θ range of 5° to 80°. Data were analyzed using MDI Jade9 and PDF-2 (release 2020), and the results were plotted using GraphPad Prism (release 8.0.2).

### Evaluation of effects of microbial dark oxygen on deep-sea microorganisms

To investigate the impact of oxygen produced by *Deferribacterota* strains DY0037 and DY0609 on the growth of deep-sea microorganisms, we designed a specialized setup. The setup consisted of an open centrifuge tube containing various media, placed inside a 500 mL anaerobic bottle. Initially, *Deferribacterota* strains DY0037 and DY0609 were cultured in the 500 mL anaerobic bottles containing 300 mL of T13 medium for two days. After this incubation period, deep-sea microorganisms were inoculated into the open centrifuge tube containing different media (*Methanolobus* sp. ZRKC1 in M1 medium (1.0 g/L NH_4_Cl, 1.0 g/L CH_3_COONa, 1.0 g/L NaHCO_3_, 1.0 g/L yeast extract, 1.0 g/L peptone, 1.0 g/L cysteine hydrochloride, 500 µL/L 0.1 % (w/v) resazurin, 10 mL/L methanol, 1.0 L filtered seawater, pH 7.0), *Erythrobacter* sp. 21-3 in 2216E (5.0 g/L peptone, 1.0 g/L yeast extract, 1.0 g/L cysteine hydrochloride, 500 µL/L 0.1 % (w/v) resazurin, 1.0 L filtered seawater, pH 7.0) or 2216E supplemented with 40 mM Na_2_S_2_O_3_). After three days of static incubation, OD_600_ was measured via the microplate reader.

To quantify zero-valent sulfur (ZVS) produced by *Erythrobacter* sp. 21-3 under anaerobic conditions in 2216E medium supplemented with 40 mM Na₂S₂O₃, we used a previously described trichloromethane-based extraction method (*58–60*). Specifically, we performed a two-step extraction, treating 2 mL of the culture twice with 5 mL of trichloromethane. The extracts were then measured using a UV–Vis spectrometer (Infinite M1000 Pro; Tecan, Männedorf, Switzerland) at 270 nm. Due to the high turbidity in the culture resulting from ZVS formation, optical density (OD_600_) measurements were unreliable for growth monitoring. Therefore, we quantified growth by colony-forming units (CFUs), using serial dilution and plating on 2216E agar, followed by a 3-day incubation.

### Detection of anaerobic manganese oxidation and nodule formation

To assess the contribution of oxygen generated by *Deferribacterota* strains DY0037 and DY0609 to manganese nodule formation, two strains were cultured in T16 medium (0.02 g/L Na_2_SO_4_, 0.05 g/L KH_2_PO_4_, 0.05 g/L Na_2_SiO_3_, 1.0 g/L NH_4_Cl, 20.0 g/L NaCl, 3.0 g/L MgCl_2_, 0.5 g/L KCl, 0.15 g/L CaCl_2_, 1.0 g/L NaNO_3_, 1.0 g/L CH_3_COONa, 1.0 g/L NaHCO_3_, 1.0 g/L cysteine hydrochloride, 500 µL/L 0.1 % (w/v) resazurin, 10 mL/L trace element solution, 1.0 L sterilized distilled water, pH 7.0) supplemented with 20 mM MnCl_2_ at 28 °C for 4 days. The resulting precipitates were collected and analyzed using TEM and field emission scanning electron microscopy (FESEM, ZEISS Gemini500, Germany). Elemental composition and content were analyzed using energy dispersive spectroscopy (EDS, Ultim Max 40, Oxford Instruments, UK) coupled with FESEM. Additionally, X-ray photoelectron spectroscopy (XPS, ESCALAB 250Xi, Thermo Fisher, USA) was employed to determine the ratios of different manganese oxidation states.

### Amplification sequencing analysis

To investigate the potential impact of oxygen produced by *Deferribacterota* strains DY0037 and DY0609 on surrounding microbial communities, we used dialysis bags containing 100 mL of T13 medium inoculated with strains DY0037 and DY0609 (experimental groups) or without bacteria (control group). These bags were placed inside 500 mL anaerobic bottles, each containing 350 mL of T13 medium and a homogenous suspension of deep-sea hydrothermal vent sediment. Three biological replicates were prepared for each condition. After a 7-day static incubation at 28 °C, cells were harvested from all bottles, and operational taxonomic units (OTUs) sequencing was performed by Novogene (Tianjin, China). Total DNAs from these samples were extracted by the CTAB/SDS method (*61*) and were diluted to 1 ng/µL with sterile water and used for PCR template. 16S rRNA genes of distinct regions (16S V3/V4) were amplified using specific primers (341F: 5’-CCTAYGGGRBGCASCAG - 3’ and 806R: 5’-GGACTACNNGGGTATCTAAT-3’) with the barcode. These PCR products were purified with a Qiagen Gel Extraction Kit (Qiagen, Germany) and prepared to construct libraries. Sequencing libraries were generated using TruSeq® DNA PCR-Free Sample Preparation Kit (Illumina, USA) following the manufacturer’s instructions. The library quality was assessed on the Qubit@ 2.0 Fluorometer (Thermo Scientific) and Agilent Bioanalyzer 2100 system. And then the library was sequenced on an Illumina NovaSeq platform and 250 bp paired-end reads were generated. Paired-end reads were merged using FLASH (V1.2.7, http://ccb.jhu.edu/software/FLASH/) (*62*), which was designed to merge paired-end reads when at least some of the reads overlap those generated from the opposite end of the same DNA fragments, and the splicing sequences were called raw tags. Quality filtering on the raw tags were performed under specific filtering conditions to obtain the high-quality clean tags (*63*) according to the QIIME (V1.9.1, http://qiime.org/scripts/split_libraries_fastq.html) quality controlled process. The tags were compared with the reference database (Silva database, https://www.arb-silva.de/) using UCHIME algorithm (UCHIME Algorithm, http://www.drive5.com/usearch/manual/uchime_algo.html) (*64*) to detect chimera sequences, and then the chimera sequences were removed (*64*). And sequences analyses were performed by Uparse software (Uparse v7.0.1001, http://drive5.com/uparse/) (*65*). Sequences with ≥97% similarity were assigned to the same OTUs. The representative sequence for each OTU was screened for further annotation. For each representative sequence, the Silva Database (release 111, 2012-07, http://www.arb-silva.de/) (*66*) was used based on Mothur algorithm to annotate taxonomic information.

### Growth assays of animals in the presence of microbial dark oxygen

To evaluate whether oxygen produced by *Deferribacterota* strains DY0037 and DY0609 can extend animal survival, we conducted survival experiments using ants and locusts. In the ant survival experiment, strains DY0037 and DY0609 were cultured for two days in 250 mL anaerobic bottles containing 200 mL of T13 medium, with the control group consisting of only T13 medium without bacterial inoculation. Five ants were placed in each 15 mL anaerobic tube, and the tube was connected to the anaerobic bottle using a blood collection needle. For the locust survival experiment, strains DY0037 and DY0609 were cultured for two days in 500 mL anaerobic bottles containing 400 mL of T13 medium, with the control group consisting of only T13 medium. Five locusts were placed in each 250 mL anaerobic bottle, which was connected to the 500 mL anaerobic bottle using a blood collection needle. All experimental procedures and equipment handling were carried out inside an anaerobic chamber. Survival time for each ant and locust was recorded.

### Gene deletion in the *Pseudomonadota* strain DY0742

To generate a *norBCD* deletion mutant of *Pseudomonadota* strain DY0742, we first amplified chromosomal fragments using primers listed in Table S1. The purified PCR products were then ligated into the suicide vector pEX18Gm, incorporating an *oriT* for conjugation. The resulting plasmid was transformed successively into *E. coli* SY327 and *E. coli* S17-1 using the CaCl_2_ method. Mating between strain DY0742 and *E. coli* S17-1 containing the plasmid was carried out at 28 °C for 48 hours. Colonies that grew on LB agar plates supplemented with chloramphenicol (Cm, 25 μg/mL) and gentamicin (Gm, 25 μg/mL) were considered single-event positive recombinant strains. Individual colonies were picked and cultured overnight at 28 °C with shaking in LB broth containing Cm (25 μg/mL) and Gm (25 μg/mL). The cultures were then diluted 1:1000 into fresh LB broth and plated on LB agar plates supplemented with 10% sucrose. After incubation for 48 hours at 28 °C, a single colony was re-streaked several times to ensure purity, followed by replication onto LB agar plates with Gm (25 μg/mL) to confirm sensitivity to gentamicin and loss of the pEX18GM vector. All double recombination mutant candidates were verified by PCR amplification and sequencing. Finally, the wild-type strain DY0742 and its *norBCD* mutant were cultured in T13 medium under both anaerobic and aerobic conditions at 28 °C for two days. The optical density at 600 nm (OD_600_) was measured using a microplate reader.

### Genomic analysis of nitrogen metabolism genes in deep-sea microorganisms

To investigate the relationship between oxygen production and nitrogen metabolism in deep-sea microorganisms, we analyzed the genomes of several currently available deep-sea pure cultures from our lab using seven databases: Pfam, GO, KEGG, COG, NR, TCDB, and Swiss-Prot. We quantified and compared the distribution of genes involved in dissimilatory nitrate reduction, denitrification, nitrogen fixation, and other nitrogen metabolic pathways in deep-sea bacterial isolates, distinguishing between those with and without oxygen production capability.

### Data availability

The full-length 16S rRNA gene sequences of deep-sea bacterial strains DY0037, DY0609, DY0563, DY0809b, DY0742, 24LQ007, and 273 have been deposited at GenBank under the accession numbers PV030186, PV030187, PQ827512, PQ827461, PQ827347, PQ827449, and KU529971, respectively. The complete genome sequences of bacterial strains DY0037, DY0609, DY0563, DY0809b, DY0742, 24LQ007, and 273 have been deposited at GenBank under the accession numbers CP179913, CP179935, CP179914, CP179936, CP179937, CP179938, and CP015641, respectively. The raw sequencing reads from the transcriptomics analyses of DY0037 and DY0609 strains cultured in T13 medium supplemented with different sodium nitrate have been deposited into the NCBI Short Read Archive (accession numbers: PRJNA1216325 and PRJNA1218497, respectively). The raw sequencing reads from the amplification sequencing analyses of strains DY0037 and DY0609 have been deposited into the NCBI Short Read Archive (accession number: PRJNA1218488).

### Statistical analysis

All data were expressed as means ± SD. Statistical analysis was performed using GraphPad Prism 8.0.2. Independent-sample *t*-test and single-sample *t*-test were conducted to determine the significance between groups. Differences of *P* < 0.05 were considered statistically significant (\**P* < 0.05, \*\**P* < 0.01, \*\*\**P* < 0.001, \*\*\*\**P* < 0.0001). Differences of *P* > 0.05 were considered not significant (ns).

## Acknowledgements

This work was funded by the NSFC Innovative Group Grant (No. 42221005), Major Research Plan of the National Natural Science Foundation (Grant No. 92351301), Science and Technology Innovation Project of Laoshan Laboratory (Grant Nos. LSKJ202203103 and 2022QNLM030004-3), Shandong Provincial Natural Science Foundation (Grant Nos. ZR2024ZD49 and ZR2021ZD28), and Taishan Scholars Program (Grant Nos. tstp20230637 and tsqn202312264). We are grateful to all scientist and crew member of ShenhaiYihao and the pilots of HOV Jiaolong during the cruise of DY83 for the sample collection. We extend special appreciation to the captain and crew of the *RV KEXUE*, as well as the FAXIAN ROV team for assistance with sample collection.

## Author contributions

RZ, CW, and CS conceived and designed the study; RZ and CW conducted most of the experiments; CW, HW, and HQ collected the samples from the deep-sea hydrothermal vents, cold seeps and seamounts; RZ, CW, and CS lead the writing of the paper; all authors contributed to and reviewed the paper.

## Notes

### Competing Interest Statement

The authors have declared no competing interest.

### Summary of Updates

We polished the writing of this paper.

