## Supplemental Table 1 for "Nitrate-driven dark oxygen production by diverse deep-sea microorganisms"

### Supplementary Methods

**Transcriptional profiling of *Deferribacterota* strains DY0037 and DY0609 cultured in T13 medium supplemented with different concentrations of sodium nitrate**

**(1) Library preparation for strand-specific transcriptome sequencing.** A total amount of 3 μg RNA per sample was used as input material for the RNA sample preparations. Sequencing libraries were generated using NEBNext® Ultra™ Directional RNA Library Prep Kit for Illumina® (NEB, USA) according to manufacturer's recommendations and index codes were added to attribute sequences to each sample. The rRNA is removed using a specialized kit that leaves the mRNA. Fragmentation was carried out using divalent cations under elevated temperature in NEBNext First Strand Synthesis Reaction Buffer (5×). First strand cDNA was synthesized using random hexamer primer and M-MuLV Reverse Transcriptase (RNaseH$^-$). Second strand cDNA synthesis was subsequently performed using DNA Polymerase I and RNase H. In the reaction buffer, dNTPs with dTTP were replaced by dUTP. Remaining overhangs were converted into blunt ends via exonuclease/polymerase activities. After adenylation of 3' ends of DNA fragments, NEBNext Adaptor with hairpin loop structure was ligated to prepare for hybridization. In order to select cDNA fragments of preferentially 150~200 bp in length, the library fragments were purified with AMPure XP system (Beckman Coulter, Beverly, USA). Then 3 μL USER Enzyme (NEB，USA) was used with size-selected, adaptor-ligated cDNA at 37 °C for 15 minutes followed by 5 minutes at 95 °C before PCR. Then PCR was performed with Phusion High-Fidelity DNA polymerase, Universal PCR primers and Index (X) Primer. At last, products were purified (AMPure XP system) and library quality was assessed on the Agilent Bioanalyzer 2100 system.

**(2) Clustering and sequencing.** The clustering of the index-coded samples was performed on a cBot Cluster Generation System using TruSeq PE Cluster Kit v3-cBot-HS (Illumia) according to the manufacturer's instructions. After cluster generation, the library preparations were sequenced on an Illumina Hiseq platform and paired-end reads were generated.

**(3) Data analysis.** Raw data (raw reads) of fastq format were firstly processed through in-house perl scripts. In this step, clean data (clean reads) were obtained by removing reads containing adapter, reads containing ploy-N and low quality reads from raw data. At the same time, Q20, Q30 and GC content the clean data were calculated. All the downstream analyses were based on the clean data with high quality. Reference genome and gene model annotation files were downloaded from genome website directly. Both building index of reference genome and aligning clean reads to reference genome were used Bowtie2-2.2.3 (*1*). HTSeq v0.6.1 was used to count the reads numbers mapped to each gene. And then FPKM of each gene was calculated based on the length of the gene and reads count mapped to this gene. FPKM, expected number of Fragments Per Kilobase of transcript sequence per Millions base pairs sequenced, considers the effect of sequencing depth and gene length for the reads count at the same time, and is currently the most commonly used method for estimating gene expression levels (*2*).

**(4) Differential expression analysis.** Differential expression analysis of two conditions/groups (two biological replicates per condition) was performed using the DESeq R package (1.20.0) (*3*). DESeq provide statistical routines for determining differential expression in digital gene expression data using a model based on the negative binomial distribution. The resulting *P*-values were adjusted using the Benjamini and Hochberg's approach for controlling the false discovery rate. Genes with an adjusted $P < 0.05$ found by DESeq were assigned as differentially expressed. (For DEGSeq without biological replicates) Prior to differential gene expression analysis, for each sequenced library, the read counts were adjusted by edgeR program package through one scaling normalized factor. Corrected *P*-value of 0.005 and $\log_2$ (Fold change) of 1 were set as the threshold for significantly differential expression.

**(5) GO and KEGG enrichment analysis of differentially expressed genes.** Gene Ontology (GO) enrichment analysis of differentially expressed genes was implemented by the GOseq R package, in which gene length bias was corrected (*4*). GO terms with corrected *P* value less than 0.05 were considered significantly enriched by differential expressed genes. KEGG is a database resource for understanding high-level functions and utilities of the biological system, such as the cell, the organism and the ecosystem, from molecular-level information, especially large-scale molecular datasets generated by genome sequencing and other high-throughput experimental technologies (http://www.genome.jp/kegg/) (*5*). We used KOBAS software to test the statistical enrichment of differential expression genes in KEGG pathways.

**Supplementary Table 1. Primers used for the deletion of *norBCD***

| Primer name | Nucleotide Sequence (5’-3’) |
|---|---|
| pEX18-GM-f | CAAACTCCATGCACAGGCAGGGTACCGAGCTCGAATTCGT |
| pEX18-GM-r | GATCACAATCTGGTGCTCTACGGGGATCCTCTAGAGTCGA |
| *norBCD*-L-f | ACGAATTCGAGCTCGGTACCCTGCCTGTGCATGGAGTTTG |
| *norBCD*-L-r | CCTCACCAGCCGCTGCTGGCGGTCGCCTCCTATTGGCTTG |
| *norBCD*-R-f | CAAGCCAATAGGAGGCGACCGCCAGCAGCGGCTGGTGAGG |
| *norBCD*-R-r | TCGACTCTAGAGGATCCCCGTAGAGCACCAGATTGTGATC |
